# Femtosecond X-ray snapshots reveal correlated displacements of specific distal atoms in a protein crystal

**DOI:** 10.1101/2024.05.29.596429

**Authors:** Viktor Ahlberg Gagnér, Maja Jensen, Torbjörn Nur Olsson, Juan Cabello Sánchez, Åsa U. J. Bengtsson, J. Carl Ekström, Zhedong Zhang, Andrius Jurgilaitis, David Kroon, Van-Thai Pham, Stefano Checchia, Hélène Coudert-Alteirac, Helena Rodilla, Jan Stake, Vitali Zhaunerchyk, Ran Friedman, Jörgen Larsson, Stefano A. Mezzasalma, Gergely Katona

## Abstract

Protein dynamics is shaped by interactions between thermal fluctuations, external forces, and molecular structures. Thermal fluctuations have high frequencies and are hence very challenging to quantify. We were able to record femtosecond X-ray diffraction snapshots and determine atomic displacements of crystalline bovine trypsin atoms either in the native steady state or in a transient state initiated by a short THz pulse, the results of which were interpreted with numerical simulations. In the absence of THz pulses, specific distal atoms exhibited correlated movements. Under the influence of THz fields, numerical simulations demonstrated a slight enhancement of displacement correlations. The experimental results revealed a contrasting response to short THz pulses, where the measured displacements were not solely determined by the atom position, but also influenced by the atomic type. These findings call for a reinterpretation of the dynamic properties inherent in folded protein structures and the physical principles behind the formation of protein assemblies. To this end, a theoretical model was developed for lattice deformations and local excitations entraining a (squeezed) coherent steady state, explaining the emergence of atom correlations. The model accounts for the magnitudes of atomic displacements and fluctuations through a balance between harmonic and anharmonic coupling forces.

---

One of life scientists’ highest goals is to predict how proteins function in real time at atomic resolution. Some proteins are short–lived in the cell and may only last a few minutes or hours, while others can remain active for weeks, months, or even years. Understanding biology in detail, from femtoseconds (fs) to at least minutes, is also required to mimic nature in artificial design. Knowledge of static three-dimensional structures and key molecular interactions alone proved ineffective in dealing with biological function, as it ultimately depends on the dynamic “character” encoded by protein molecules in continuous time and space^1^. Atomic displacements (or shifts) in a folded protein core must be delicately balanced at the time scales relevant to proteins. How do thermal fluctuations at organism-specific temperatures, ranging from arctic organisms to extremophiles thriving at hot hydrothermal vents, lead to convergence towards protein structures rather than divergence to a disordered state?

To better understand the occurrence of ordered structures in biological or crystallographic contexts, the focus is often placed on the constraints imposed by physical space and chemistry. For instance, the presence of 65 Sohncke space groups in protein crystals can be explained by the necessity of maintaining translational symmetry. Similarly, biological structures are influenced by the rigid geometry of covalent bonds in functional groups, as well as the optimal coordination of hydrogen bonds, metal-ligand interactions, etc. Such considerations frequently ignore that an optimal geometry can be achieved in a disordered state (in a manner that minimizes the energy). A protein can form better hydrogen bonding with the solvent and assume a less strained, extended geometry, when it is not in a folded state. In fact, many proteins exist in a disordered state either permanently, as in intrinsically disordered proteins, or temporarily. An example for the latter situation is a folded protein that loses its structure outside its optimal temperature, or one that is unstructured without a partner protein before they form a complex.

One way to study protein folding is to use a statistical or knowledge-driven approach, e.g. by focusing on eliminating impossible structures, such as those endowed with significant steric clashes. A more captivating investigation can revolve around understanding the underlying forces driving the formation of ordered structures. Such investigations can reveal factors that contribute to the formation of specific point and space groups, protein folds and conformations, compared to others.

For biological macromolecules the most successful method for predicting protein structures relies on deep learning. However, this does not afford a fundamental understanding of protein structures, protein assemblies or their crystals. The most commonly used model, to explain how proteins adapt to their 3D structures, is based on energy funnels with deep minima^2^. Accordingly, any conformation is posited to correspond to a long-term steady state of a (relative) energy minimum. This explanation, unfortunately, is difficult to test experimentally. The funnel theory implies an enormously complex conformational energy surface, the deepest valley of which specifying the native state, with huge numbers of local minima (approximately 10^100^ per protein domain^3^) connected by saddles. It cannot be modelled exhaustively, while insights from the funnel approach to folding are insufficient to improve computer searches^4^. Further shortcomings of the funnel model are that it does not account for thermal fluctuations in ambient temperatures, for allosteric folding and for proteins adopting multiple states.

In the analysis of protein structure and dynamics, including that of macromolecular assemblies and crystals, an alternative viewpoint that is built on mode-locked oscillating dipoles comes handy. According to this view, dipoles synchronize their oscillation phase and attain regular oscillation direction. To make this concept useful, it is essential to pinpoint which elements exhibit dipole behaviour. We can start with excluding groups of elements: not all atoms in a biological macromolecule can simultaneously participate in the same mode-locked dynamics, as identical macromolecules are rarely detected to be oriented exactly in the same direction in protein assemblies or crystals. Instead, we observe a predominance of stable structures that exhibit rotational invariance, intramolecular and intermolecular symmetry. Rotational invariance and symmetric structures are often highlighted for their utility, leading to discussions to depict them as products of evolution^5-7^. Symmetric structures can be useful because genetic information can be compacted to encode only a single subunit which self-assembles with itself to form a large object. A symmetric assembly can be more resilient against deformations^8^ and denaturation^9^. The assembly can gain new allosteric regulation that a single subunit does not have, for example the cooperative action of haemoglobin subunits facilitates oxygen take up and release^9^.

It is worth mentioning that rotationally symmetric structures can also occur without any apparent utility. The Protein Data Bank contains numerous crystal structures with rotational symmetry. In the following discussion we are concerned by a subset of these, where the rotational symmetry exists in the biological unit, rather than merely in the crystal. Taking a utility-independent perspective on the physical mechanism leading to the formation of rotational invariance involves the building up of a restricted and homogeneous subset of atoms that establish a symmetry axis through their collective wavevector. In that case, the remaining atoms of the protein would be either subject to incoherent dynamics with greater degrees of freedom or partake in separate coherent processes. Mode locking of distal sites provides a fresh outlook on the allosteric behaviour of proteins, which appears in turn to be a general property of folded structures^10^. Current models to explain allostery describe the functional coordination in a large protein domain^11^ by an elastic network of nodes with local spring interactions that propagate signals between distal sites in response to mechanical (network) deformations.

Experimental studies of atom correlations are challenging. Coherent quasi–elastic neutron scattering (QENS) is an important source of evidence for collective shifts of protein atoms^12^. In protein solutions, a picosecond (ps) dynamics develops in–phase up to nanometer (nm) scales. Because QENS does not provide atomic resolution, determining which atoms move in-phase remains a tough task. Due to the large number of degrees of freedom in a protein, the collectivity of ps motions may be limited to individual molecules and the (proximally) perturbed water network. Terahertz (THz) absorption spectroscopy becomes useful in this context. With increasing protein concentrations, hydration shells begin to overlap, resulting in a non–monotonic change in the concentration dependence of THz absorption spectra^13^. Long–range EM forces were detected by THz spectroscopy between protein molecules in solution under out-of-equilibrium conditions^14,15^, which may have implications for their macroscopic behavior. Anisotropic THz microscopy revealed that coherent and collective oscillatory regimes take place in protein crystals, where variations of spectroscopic signals were recorded against the crystal orientation relative to the THz field polarization^16-19^. As compared to static steady states, coherent or collective ordered oscillations can encode and convey more information, yet their range and nature in proteins and protein crystals are largely unknown.

In protein solutions, a steady state superposition of solvent–slaved and non–slaved motions can take place in large conformational domains^20,21^ on slower time scales. Solvent effects, however, could be also important at ultrafast time scales (fs) and with small amplitude fluctuations. Large–scale conformational transitions associated with substrate binding in enzymes^22^ and light activation^23-25^ are not instantaneous, but trigger fast fluctuations that maintain a steady state until the reaction advances. Finally, low–frequency (THz) vibrations were recently suggested to cover an important role in the function and dynamics of biomolecular structures (e.g.^26^). Correlated fluctuations were proposed to go into a resonant state and form a coherent pathway of allosteric propagation in G-protein coupled receptors^27^. As initially surmised by Fröhlich^28^, they can give rise to a non-thermal steady–state, whose energy is mostly distributed to long-lived excited phonon modes of the lowest frequencies. Slow vibrational modes can be elicited by the environment (thermal bath, *kT* ≈ 200 cm^-1^) and are known to partake in thermally activated barrier crossings. Utilizing the coherent phenomena similar to Fröhlich’s condensation would have many implications in biology and medicine. Consider biochemical reaction rates as an example; these may vary exponentially with the energy in the lowest mode, producing dramatic changes in comparison with scenarios in which the input energy gets randomly distributed^29^.

In this work, we aimed to visualize collective movements in protein crystals and provide a model description in terms of coherent waves. Experimentally, the ps dynamics of atomic shifts in bovine trypsin was studied by fs–snapshots of crystal states at atomic resolution^30-32^. One of the primary questions addressed here is whether there is evidence for THz-induced elastic deformations of the protein unit cell before thermalisation completes. Having addressed this issue, we probed a number of long-range dynamic similarities in the behaviour of distal atom displacements^33^ which, although being elastic in nature, cannot be framed into a classical theory for the movement or the deformation of a rigid or elastic body (such as in ordinary gel or polymer networks^34^) and cannot arise by chance. MD simulations of the crystal, subject to electric fields that matched the experiment, were also run and analysed. Molecular dynamics (MD) simulations may have the unbeatable edge to (partly) go in atomic detail, but they should be seen more as experiments than comprehensive theories. Consequently, we adopted a theoretical approach decomposing our problem into dual quasiparticle dynamics, where shifts and fluctuations of phonon lattice modes interact with vibrational excitations of a more localized nature in coherence conditions.

## Proteins have a mechanism for extending their assembly symmetry over macroscopic distances

A pertinent issue highlighted here is the non-coincidental alignment between symmetry elements present in biological assemblies and those observed in their crystal structures, as this phenomenon lacks an adequate explanation. It is widely believed that the formation of biological assemblies stems from short-range interaction patterns among their subunits, while the creation of crystal structures involves an additional set of crystal contacts which dictate the positioning and orientation of symmetry elements. These two sets of symmetry axes, one related to biological assembly and the other to crystal formation, do not necessarily align. In fact, there are cases of universal discrepancies, such as when the 5- and 7-fold symmetry of a protein assembly clashes with the translational symmetry of a crystal lattice. For example, this occurs in PDB ID 3T30 of human nucleoplasmin (Npm2)^35^ and PDB ID 3JBT of Apaf-1 apoptosome^36^. However, it is noteworthy that a significant portion of protein crystal structures do exhibit assembly symmetry axes that align precisely with their crystal symmetry axes, despite the expectation that these two sets of short-range interactions would operate autonomously (Table 1).

**Table 1:** Survey of human homooligomeric protein assemblies in the PDB. The table values represent the number of unique Uniprot entries. The content of the asymmetric unit (ASU) is analysed: for each listed assembly symmetry, the ASU containing exactly one chain (Alternative 1) and exact multiples of chains corresponding to that assembly symmetry (Alternative 2) in the ASU, is selected.

| Assembly symmetry | ONLY crystals containing 1 chain/ASU Alternative 1 | BOTH crystals 1 chain/ASU AND more than 1 chains/ASU Alternative 1 & 2 | ONLY crystals containing more than 1 chains/ASU Alternative 2 | Fraction of proteins with equivalent assembly and crystallographic symmetry (Alternative 1) |
| --- | --- | --- | --- | --- |
| C2 | 397 | 344 | 1219 | 38% |
| C3 | 49 | 29 | 86 | 48% |
| C4 | 17 | 5 | 13 | 63% |
| D2 | 48 | 18 | 156 | 30% |
| D3 | 24 | 2 | 41 | 39% |
| D4 | 8 | 2 | 6 | 63% |
| Any of these assembly symmetries | 543 | 400 | 1521 | 38% |

We evaluate the likelihood of aligning assembly symmetry to crystallographic axis by considering their orientation. A straightforward estimate for the probability of misalignment (*p* = Δ Ω/4π) can be derived by using the concept of solid angle (*dΩ* = *sinФdθdФ*), applied, for simplicity, to a unit misalignment vector. Assume that any polar deviation of up to 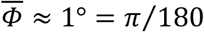 rad between crystallographic and assembly axes is imperceptible in experimental measurements. Then the expression 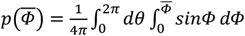 provides a minimal formula for the desired probability, θ denoting the azimuth angle with values uniformly distributed on the circle (Figure S1).

For small angles, the approximation 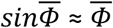 holds, finally resulting in 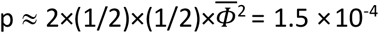. As oppositely oriented vectors yield parallel axes (say, 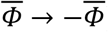), we have doubled the cumulative probability. Alignment of nearly parallel symmetry axis positions imposes additional constraints, but an exact probabilistic evaluation lies beyond the scope of this analysis and is not provided here. Nevertheless, the occurrence of such an alignment would be highly unexpected if crystal contacts and biological interfaces were solely responsible for determining these two axes separately.

A survey of the PDB database defies this expectation. 38% of human proteins crystallized and modeled with only one chain (Table 1), despite being attributed with assembly symmetry of C2, C3, C4, D2, D3 or D4. This suggests an alignment between solution assembly and crystallographic symmetry. The fraction of crystal structures that can be described with one chain increases as the symmetry of structures is enhanced both in cyclic and dihedral series. The survey encompassed 2464 unique human protein chains forming purely homooligomeric assemblies, and this can be contrasted with the 20030 protein chains in the human proteome, their fraction that do not belong to Intrinsically Disordered Proteins (IDPs), totaling 13564 proteins^37^ or the coverage of human protein chains in the PDB (7085)^38^. Among the human protein chains (at least some fragment of them) documented in the PDB with their Uniprot ID, 35% have the capability to form one of these simple homooligomeric assemblies (excluding symmetries higher than 4-fold, cubic and helical ones, and symmetries present only in heterooligomers), while 13% of them can extend their assembly symmetry to macroscopic distances. This significant fraction directly exposes a physical mechanism that links a group of entities defining the symmetry axis. In numerous cases, crystallographic and non-crystallographic symmetry axes diverge, allowing for different and independent sets of elements defining the axes. However, there is no need to introduce a separate physical mechanism to account for crystallographic and non-crystallographic (including assembly) symmetry; the action of the same mechanism on different and readily available elements suffices. In summary, biological symmetry is prevalent in human proteins, and a significant portion of protein assembly crystal structures cannot be adequately explained by the mechanism of short-range assembly and crystal contacts alone. In this study, we tackled the elements responsible for the long-range organization of protein molecules and propose a mechanism that enables the formation of symmetric structures.

## Experimental THz measurements and pulse timing reveal no changes that affects the crystal unit cell

Remarkably, single-cycle THz pulses did not modify the integrity of protein crystals even after many thousands of repetitions. The experimental station and pulse timing are outlined in Figure 1. Even-numbered images of diffraction intensity (labelled by i, *DI*_*THz*,*i*_) were measured with 50 ps delay from the application of a THz pump pulse. Odd-numbered ones (*DI*_*off*,*i*_) were recorded without initial THz pulses. To better account for the variability of results, original diffraction images (one per X-ray pulse, roughly 134,400 in total) were regrouped into *n* (=74) different combinations in each dataset, *DI*_*THz*,*i*_ and *DI*_*off*,*i*_. Only a randomly selected half of THz and off images were employed before summation, avoiding the replacement of the same diffraction image in the resampling procedure at the only expense of a small loss of data precision. Resampled data allowed to get unique amplitudes for the structure factors, *F*_*THz*,*n*_ and *F*_*off*,*n*_ respectively from 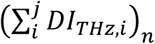 and 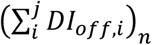. The total number of images that were kept in both sums to j = 25 at each resampling rotation. An automated refinement pipeline returned two sets of crystallographic models, *M*_*THz*,*n*_ and *M*_*off*,*n*_, from which a deformation field, generally arising as a model difference, *M*_*THz*,*n*_ −*M*_*off*,*n*_, was finally defined. Its character depended on the tensor rank of the quantities compared, i.e. it was vectorial for the displacement of mean positions (bold notation) or scalar for B-factor changes between THz on and off states (i.e. *B*_*THz*,*n*_ − *B*_*off*,*n*_). By the notations *M*_*a*,*THz*,*n*_, *M*_*b*,*THz*,*n*_ or *M*_*a*,*off*,*n*_, *M*_*b*,*off*,*n*_, any such field can be more specifically tailored to address pairs of atoms, *a* and *b*, picked up from the models *M*_*THz*,*n*_ and *M*_*off*,*n*_. The resampling process, data reduction and modelling strategy are illustrated in Figure 1. Table S1 reports the averages and standard deviations of the quality indicators for structure factors and models, *F*_*THz*,*n*_, *F*_*off*,*n*_, *M*_*THz*,*n*_, *M*_*off*,*n*_, which display negligible differences. The standard deviations of cell parameters hide the large correlation between errors on odd and even datasets, with their differences staying basically constant. Owing to a number of interruptions during the acquisition, diffraction data were processed in multiple rotation wedges, indexed independently, and it turned out that introducing additional (yet unnecessary) orientation–dependent modelling errors between wedges increased the uncertainty on cell parameters. Experimental and modelling errors appeared to be highly correlated only when one rotation wedge was employed (Table 2). This trend confirms our earlier studies^33,39^, where the positive correlation in experimental errors was helpful to improve single wavelength diffraction phasing^40^. The solvent content of the crystals and the potential for thermal effects to perturb the conformational equilibrium in trypsin are discussed in SI-1a and SI-1b.

**Table 2:** Data reduction statistics (first wedge). The mean and standard deviation are displayed. The values in parenthesis refer to the reflections in the highest resolution bin.

|  | **F<sub>off,n</sub>** | **F<sub>THz,n</sub>** |
| --- | --- | --- |
| Resolution range (Å) | 6.44 - 1.52 (1.56 - 1.52) | 6.44 - 1.52 (1.56 - 1.52) |
| Space group | P2<sub>1</sub>2<sub>1</sub>2<sub>1</sub> | P2<sub>1</sub>2<sub>1</sub>2<sub>1</sub> |
| Unit cell A length (Å) | 55.250±0.003 | 55.251±0.004 |
| Unit cell B length (Å) | 59.021±0.004 | 59.023±0.006 |
| Unit cell C length (Å) | 67.963±0.003 | 67.966±0.004 |
| Total number of reflections | 40837±39 (1139±5) | 40791±52 (1140±7) |
| Number of unique reflections | 23645±22 (810±4) | 23624±27 (810±7) |
| Multiplicity | 1.7±0.0 (1.4±0.0) | 1.7±0.0 (1.4±0.0) |
| Completeness (%) | 57.7±0.1 (28.7±0.1) | 57.7±0.1 (28.8±0.2) |
| CC<sub>1/2</sub> (%) | 99.0±0.0 (48.8±4.2) | 99.0±0.0 (51.0±3.7) |
| <I/σ(I)> | 5.76±0.04 (1.12±0.03) | 5.77±0.05 (1.07±0.02) |
| R<sub>merge</sub> (%) | 8.3±0.1 (49.9±2.1) | 8.2±0.1 (51.8±2.4) |
| R<sub>meas</sub> (%) | 10.8±0.1 (70.6±3.0) | 10.7±0.1 (73.3±3.4) |
| Wilson B-factor (Å<sup>2</sup>) | 18.707±0.084 | 18.500±0.082 |
| Beam divergence (°) | 1.109±0.007 | 1.105±0.011 |
| Mosaicity (°) | 0.107±0.003 | 0.108±0.004 |

**Figure 1:**
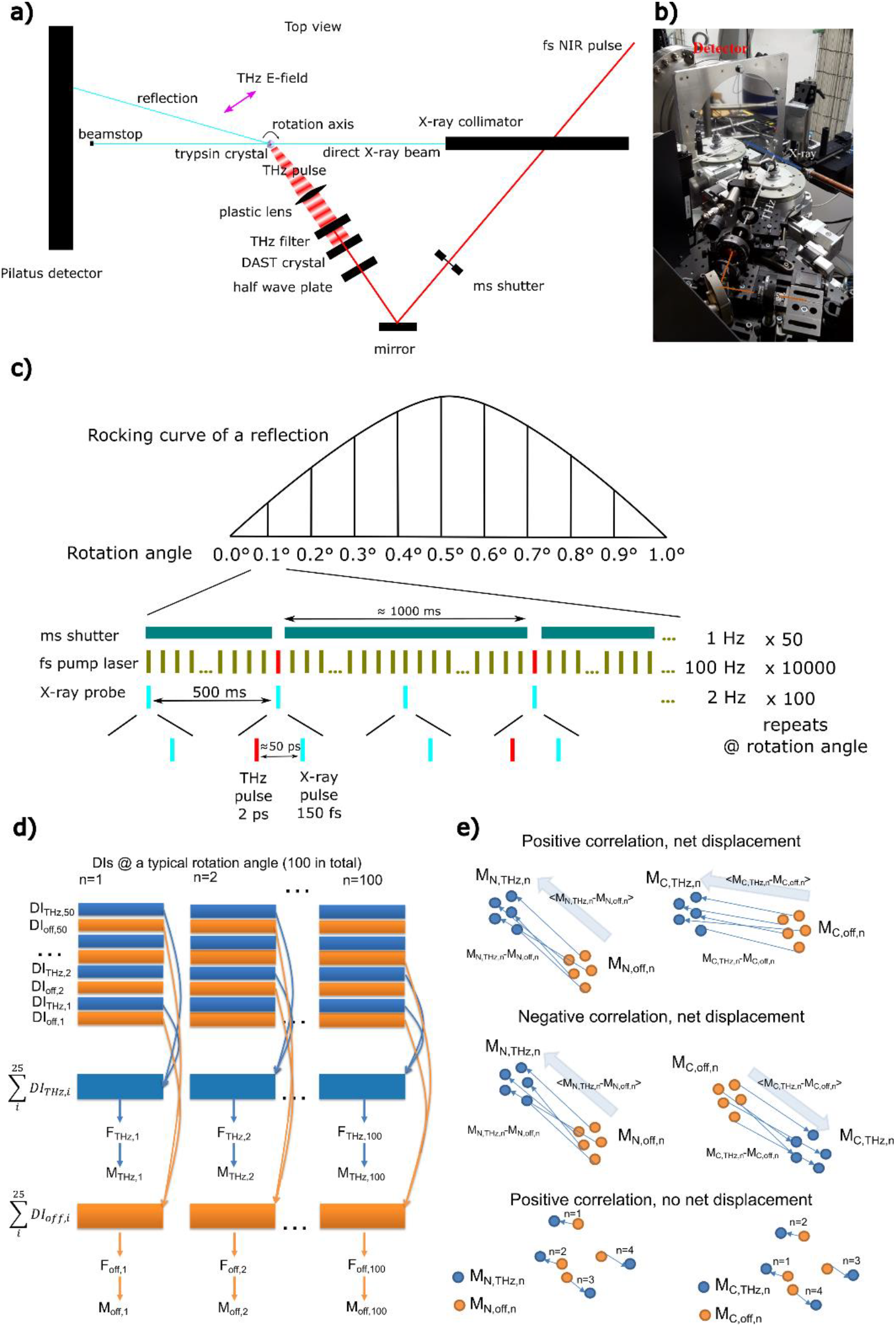
Experimental station and timing scheme. a) X-ray beampaths with two alternative probabilites are depicted in cyan colour. The red line indicates the beampath of NIR pump laser. Optical rectification on the DAST crystal generates the THz pulse (shown by a wave pattern with macroscopic wavelength). b) A photograph of the experimental station with beampaths indicated. c) A timing scheme of the instruments. d) A scheme of resampling, data reduction and modelling. Note that generating structure factor amplitudes and structural models require diffraction images (DIs) from multiple rotation angles but, for the sake of clarity, only resampling of diffraction images at a single rotation is shown here. e) A scheme of the statistical relationship for the resampled position of atom pairs. Yellow and blue indicate THz_off_ and THz_on_ states (or positions), respectively.

In summary, THz pulses sent to bovine trypsin crystals did not induce elastic deformations (compression/expansion) of unit cell parameters larger than 0.5 pm. From uncertainty estimates derived from resampling, we could only detect some subtle perturbation of atomic distributions at the local level and in the protein structure.

## Average displacement vectors for covalent atoms under irradiation frequently deviate from their control values

Figure 2 displays the average deformation fields ⟨*M*_*THz*,*n*_ − *M*_*off*,*n*_⟩ on the main chain amide nitrogen and carbonyl carbon positions. Pairs of these atomic species are covalently bonded in a polypeptide. The ribbon representation of the molecule corresponds to one of the asymmetric units seen in the Cartesian (x-z) projection. In this orthorhombic system, symmetry-equivalent atoms are typically associated with four mean displacement vectors, two of which share the same unit vector. The magnetic polarization vector is indicated by a purple arrow. Deformation fields ⟨*M*_*THz*,*n*_ − *M*_*off*,*n*_⟩, defined on amide nitrogen and carbonyl carbon atoms, vary in their directions substantially even between covalently bonded pairs. A parallel alignment in vectors ⟨*M*_*THz*,*n*_ − *M*_*off*,*n*_⟩ is also observed e.g. on Cβ carbon (CB) atoms of Ser-195 and His-57 (Figure 2). These two atoms were previously shown to have a surprisingly similar distribution of steady-state shifts^33,41,42^ and now we show that this similarity can stem from a synchronized swarm response of specific carbons to a perturbation on the ps time scale.

**Figure 2:**
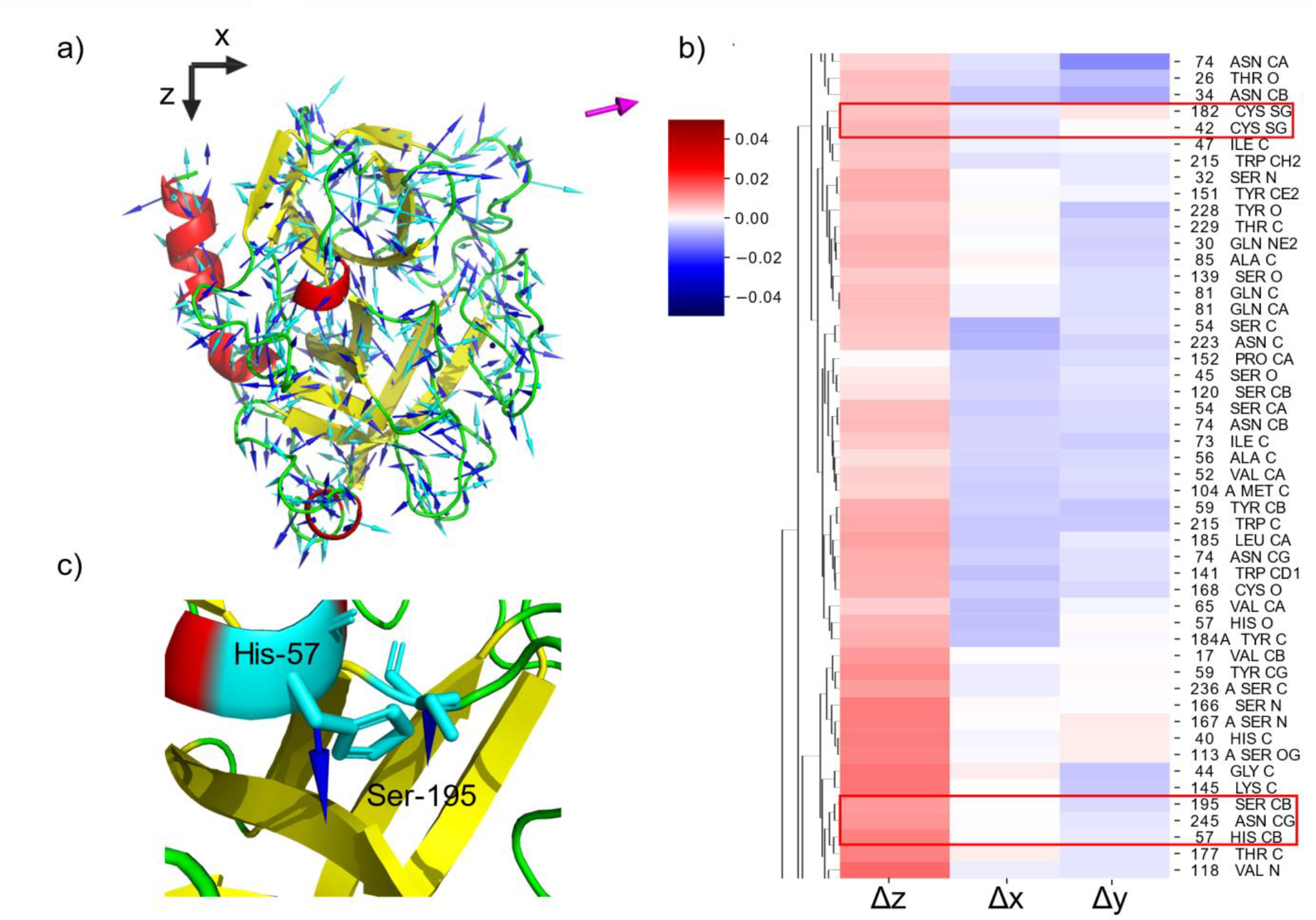
Average atom displacement directions. a) Average displacement of main chain amide nitrogen (blue arrows) and carbonyl carbon (cyan arrows) atoms upon THz excitation. The magenta arrow specifies the magnetic polarization vector of the THz pulse. Red, yellow and green colors illustrate α-helical, β sheet and loop secondary structures, respectively. Even bonded pairs of amide nitrogen and carbonyl carbon atoms often displace in different directions and with different magnitudes. b) Hierarchical clustering of deformation field vectors ⟨*M*_*THz*,*n*_ − *M*_*off*,*n*_⟩ mapped on individual atoms. For greater clarity, only 80 of total 1629 atoms are shown. Highlighted clusters show highly similar deformations of spatially separated atoms, including Cβ of Ser-195 and His-57. c) A parallel alignment of deformation fields on Cβ of Ser-195 and His-57 shown with *blue* arrows.

In addition to these atomic types, the similarity of (average) individual displacement directions rarely follows the covalent connectivity of atoms. Using hierarchical clustering, we can examine which atoms are characterized by a similar mean field, ⟨*M*_*THz*,*n*_ − *M*_*off*,*n*_⟩. Figure 2 shows a representative part of a dendrogram where, out of 80 atoms, only Gln-81 CA-C and Asn-74 CA-CB-CG correspond to covalently bonded atoms. *The deformation field is often similar on atoms that are sequentially and spatially distant from each other*. Clusters containing atoms with close chemical and functional properties are quite common, such as the sulfurs of Cys-42 and Cys-182 that are not part of the same disulfide bridge. *This is a clear sign that atoms do not respond to the THz field as part of a rigid group in an elastic scaffold*. On the other hand, the enrichment of certain atom types in the highlighted clusters points out that chemical properties govern the atomic response (Figure S2). This branch of the dendrogram reveals an enrichment of highly similar deformation of β carbon (CB), α carbon (CA), carbonyl oxygen (O), and amide nitrogen (N) atoms (Figure S2).

## Distal atom pairs exhibit a spontaneous displacement correlation, disrupted by the THz pulse

To examine the potential influence of THz-periodic electric fields (**E**), MD simulations were performed in the presence of different maximum field strengths (0-350 MV/m). Repeated calculations with randomized starting protein orientation relative to the **E-**field polarization displayed a nearly monotonic increasing behaviour of the solvent accessible surface area, and a decreasing number of H– bonds as a function of increasing strength (Figure 3a). Alongside, the magnitude of molecular dipole moments shows a jagged increasing trend (Figure 3b). These predictions emphasize the disruptive effect of the THz field on the protein structure. Since the entire molecule (and its dipole moment) does not have enough time to tumble and follow the variable **E**-vector, certain molecular domains have to reorient themselves. This leads to H–bond breakage, structural disruption, and slight unfolding with increasingly accessible surface areas. As the feedback of charge movements does not perturb the simulated fields, entrainment between atomic/molecular oscillators is not expected here.

**Figure 3:**
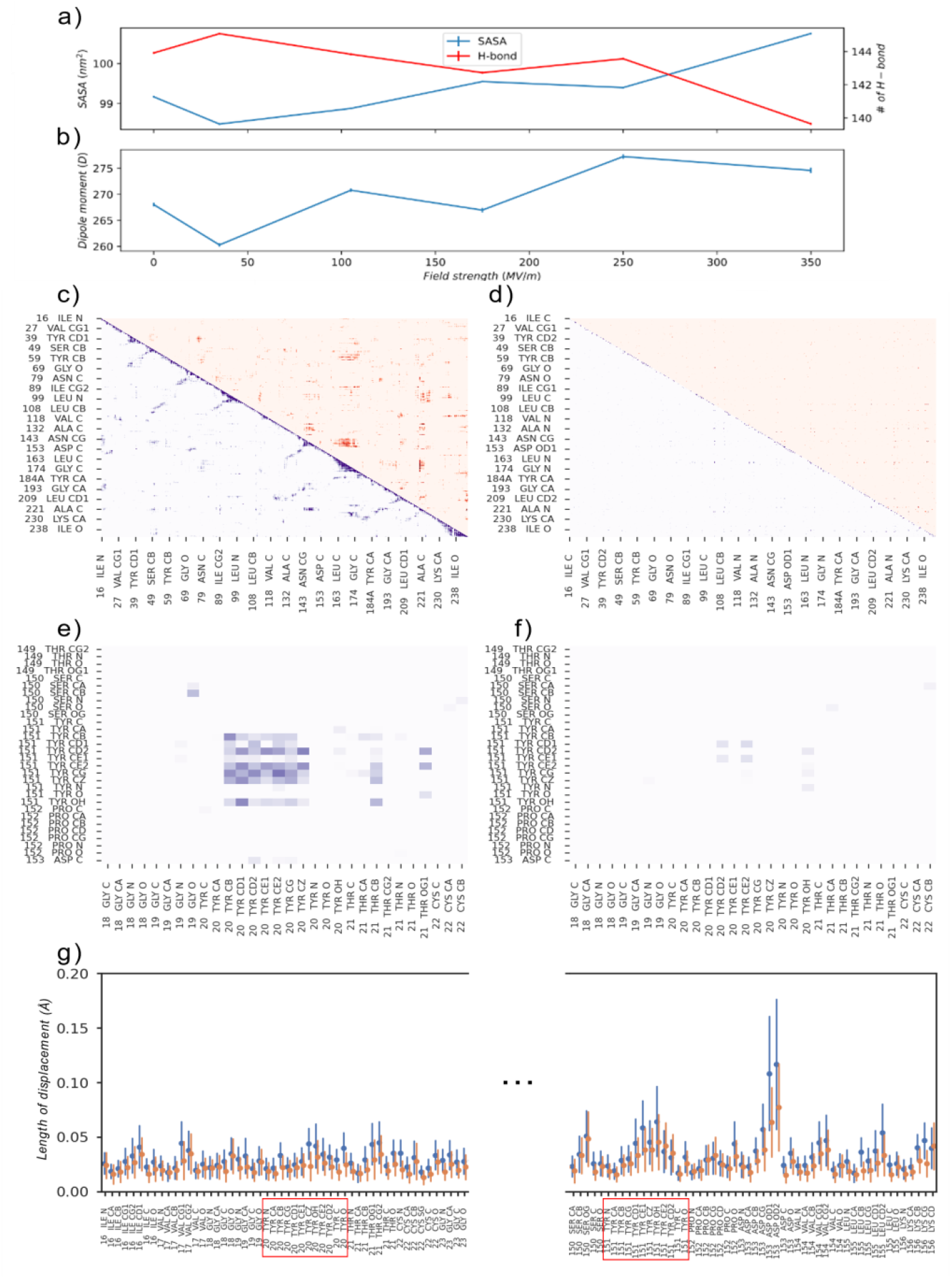
Correlation of atom displacements in MD trajectories and experimentally determined structures. a) Average and confidence intervals (CI = 0.95) of solvent accessible surface area (SASA) and number of hydrogen (H–) bonds in simulations performed in the presence of periodic **E**-field of different strengths. b) Average and confidence intervals (CI = 0.95) of dipole moment magnitude in the same set of simulations. c) Displacement correlation matrix from the MD simulation with the strongest periodic electric field (**E**). The lower and upper triangles of correlation matrices represent positive (light to dark blue: 0.2 to 0.5 or higher) and negative (light to dark red: -0.2 to -0.5 or lower) correlations, respectively. d) *corr*(*M*_*a*,*THz*,*n*_ − *M*_*a*,*off*,*n*_; *M*_*b*,*THz*,*n*_ − *M*_*b*,*off*,*n*_) of the experimental model. The correlation coefficients were calculated over the 74 resampled dataset pairs. Lower and upper triangles of correlation matrices represent positive (light to dark blue: 0.2 to 0.5 or higher) and negative (light to dark red: -0.2 to -0.5 or lower) correlations, respectively. The positive correlation between side chains of Tyr-20 and Tyr-151 are highlighted in e) *corr*(*M*_*a*,*off*,*n*_ − *M*_*a*,*off*,*m*_; *M*_*b*,*off*,*n*_ − *M*_*b*,*off*,*m*_) (*n* ≠ *m*) and f) *corr*(*M*_*a*,*THz*,*n*_ − *M*_*a*,*off*,*n*_; *M*_*b*,*THz*,*n*_ − *M*_*b*,*off*,*n*_). g) The deformation intensity is mapped on atoms of amino acid residues 16-23, including Tyr-20. The region displayed is 150-155 with Tyr-151 included. The distribution of ‖*M*_*THz*,*n*_ − *M*_*off*,*n*_‖ is shown in *blue*, while the steady state variation of ‖*M*_*off*,*n*_ − *M*_*off*,*m*_‖ (*n* ≠ *m*) is shown in *orange*.

In the experiments, every diffraction image relates to an ultrasharp snapshot of individual atoms with only 150 fs exposure time. If such snapshots were displacing from the average positions in a correlated fashion, the sum of their images would not erase the correlations. Moreover, covariance and correlation of two observables at different time points is not sensitive to the sampling order of measurement pairs. As time series can be arbitrarily rearranged without altering correlation, we can equally use well-sampled stroboscopic experiments to estimate correlation between any two observables. With this in mind, we generated correlation matrices of the form *corr*(*M*_*a*,*THz*,*n*_ − *M*_*a*,*off*,*n*_; *M*_*b*,*THz*,*n*_ − *M*_*b*,*off*,*n*_) and *corr*(*M*_*a*,*off*,*n*_ − *M*_*a*,*off*,*m*_; *M*_*b*,*off*,*n*_ − *M*_*b*,*off*,*m*_) (*n* ≠ *m*), where the semicolon demarcates the statistical deformation fields to be confronted.

Resampling crystallographic data permitted to examine the shift correlation between arbitrary atom pairs, and confront it with the results from MD simulations. We compared correlation matrices of two MD trajectories initiated from the same point with the simulation of a periodic, polarized **E**-field. Both correlation matrices show strong positive correlation close to the diagonal (Figure 3c shows a simulation performed with the strongest **E**-field). Positively correlated atom pairs are occasionally arranged in an anti-diagonal direction, corresponding to antiparallel β-sheets restrained through direct H–bonding in the simulation. A regular pattern of H–bonds is present between main chain atoms in antiparallel β-sheets. While static connectivity is a defining characteristic of elastic networks, molecular dynamics (MD) simulations continuously break and form connections. However, in well-folded protein domains, the linkage network is typically dense enough that both approaches yield similar predictions ^43^. The semi-stable connectedness among antiparallel sheets means that sequentially distal amino acid residues can be H–bonded, resulting into positively correlated displacements in anti-diagonal directions of the correlation matrix.

Negatively correlated motions appear farther away from the diagonal and usually relate to specific type of repetitive elastic deformations (twists, compressions, etc.) dictated by the protein shape and its internal connectivity. The effect of the **E**-field on correlations is not immediately obvious, but there emerges a significant increase of positively correlated atom pairs when the EM field is applied in multiple simulations (Figure S3). We can expect the ever-present electro-dynamic fields in biological matter to correlate differently charged protein atoms, albeit such changes should be small even at very large field strengths. Electrodynamics cannot be switched off in experiments as it is done in simulations, but we can introduce a momentary THz perturbation and evaluate its effect on atom-atom displacement correlations before a steady-state is restored. This experiment, however, exhibits less positive correlation along the diagonal (Figure 3d). A slightly elevated correlation is observed between, but not beyond, atoms of the same residue, and positive correlations belonging to antiparallel β-sheets are missing. Besides, strong positive correlations arise far from the diagonal between specific atomic groups.

A positive correlation between Tyr-20 and Tyr-151 side chains is highlighted in Figure 3e and 3f. Although their Cα atoms are 13 Å apart, their side chains displace in concert regardless of the network of interleaving amino acid residues. This is undoubtedly surprising if atoms in protein molecules vibrate independently, but is predicted by models that incorporate coherent or correlated oscillations in/near a steady state. High-field THz pulses appear to temporarily reduce the positive correlation to a lower level: 727796 atom pairs experience a smaller correlated displacement. Correlation is only enhanced in 599024 pairs a short delay after the single-cycle THz pulse. To explain such observations, the given atomic groups should exhibit correlated shifts in the crystal lattice prior to THz irradiation, and there is a need to investigate on how a structural steady state dynamics can be established at the level of individual molecules and across the crystal lattice. If selected atom pairs are always positively correlated in their movement, their average position will show finite correlations within many molecules, even though equivalent pairs located elsewhere are not acting in concert. As the change in average position should be negligible when millions of molecules are involved, measurable variations necessitate the synchronization of a large fraction of atomic constituents. To study this, we followed on deformation intensity distributions from the experiments.

## Deformation intensities of selected atoms are substantially larger in irradiated proteins

Figure 3g shows the deformation intensity distributions ‖*M*_*THz*,*n*_ − *M*_*off*,*n*_‖ in two regions of amino acid residues (16-22 and 150-155) that include the previously discussed Tyr-20 and Tyr-151. They can be compared to ‖*M*_*off*,*n*_ − *M*_*off*,*m*_‖), corresponding to the steady state variation in models unaffected by THz radiation. These two distributions are similar for a large number of atoms, which indicates that the latter norm is a good statistical descriptor for atoms which are not meaningfully perturbed by the THz source. There is a number of irradiated atoms with an increased deformation intensity, including Tyr-20 CZ, O, and OH. B-factor differences *B*_*THz*,*n*_ − *B*_*off*,*n*_ in Figure S4 highlight that predominant changes occur in the positive direction (Ile-16 N, C, CA, CB and CG2; Tyr-20 CE2; Ala-24 O; Tyr-29 O, C and CD2; Val-154 O; Lys-159 O and C; Ala-160 N, CA and CB; Ile-162 C and O) and, only to some extent, in that negative one (Gly-18 O; Tyr-20 O, Ser-150 CB and OG; Tyr-151 CB, CZ and OH; Val-154 CA; Leu-155 C and CA; Lys-159 N). Higher or lower correlations between distally localized directions in Figure 3e and 3f do not necessarily bring to an increased deformation intensity in Figure 3g. A strong correlation between atoms *a* and *b* can even arise when ‖⟨*M*_*a*,*THz*,*n*_ − *M*_*a*,*off*,*n*_⟩‖ = 0, provided that *a* and *b* deform in synchrony.

## Unifying long–range and short–range excitations

For explaining the high synchronization observed between protein atoms concerning the magnitude of their displacements, we formulate two out-of-equilibrium models. Such descriptions presuppose that some steady state is approached, as e.g. when the input energy rate is above a critical threshold or the system itself shows a concerted dynamics due to coupling interactions between its constituents. The theories of Fröhlich’s condensation^28^ and Davydov’s soliton^44^ provide formal structures suitable to this aim. The former predicts non–thermal long–range phonon excitations in their lowest vibrational mode^45^. In Wu–Austin’s Hamiltonian interpretation of it, ladder operators are assigned to cell–bath and heat–bath bosonic quasiparticles in a building up of molecular order^46^. The latter theory describes the longitudinal order propagation and energy transport across H–bonded spines of *α*-helix chains^47^. The interaction between some bond stretching (e.g. C=O) and the lattice deformation of H–bonds can trigger electrosolitons of amino acids travelling through the biological matter without changing shape^48^.

To combine such formalisms in a comphehensive framework, we concentrate on a minimal second quantisation model of coherence. First, the original Fröhlich’s equations for phonon occupation numbers are rearranged into a prototype dynamic system (SI–4a) and are discussed thermodynamically near (or at) the marginal onset of stability (SI–4b). It is shown that, at a high enough energy redistribution, the motion can approach/settle in an oscillatory steady state. Thereafter, we generally limit the dynamic picture to two protein excitations: intramolecular vibrations and intermolecular deformations (Figure S5a). It is proven (vide infra) that a formal structure similar to Fröhlich’s and Davydov’s theories, tailored to THz oscillators, can capture some key characteristics of displacements and fluctuations (B-factors) of protein atoms. We surmise that local vibrations couple with phonons along lattice pathways, leading to a coherent behaviour of atomic shifts. A good agreement with the experiments is achieved by adopting the typical quantities of a lattice of H–bonds coupling anharmonically with other dipole moments, as in soliton models. However, absorption low– frequencies of amino acids range overall in ∼ (0.3 − 6.2) THz^49^ and, although the predicted magnitudes are correct, the precise microscopic origin of THz coupling (i.e. which specific structural constituents, bonds, etc. are involved) remains unassigned. We will therefore stick to the main notations used in the literature, and conventionally refer to excitations of a more local nature (e.g. dipolar, at the level of amino acid residues, etc.) interacting with large–scale phononic quasiparticles (lattice deformations).

## Atom displacements in a coherent state stem from a balance between harmonic and anharmonic forces

*The starting Hamiltonian is a sum of local (a*_*k*_*), lattice (b*_*q*_*) and coupling* (*a*_*k*_*b*_*q*_) *contributions:*

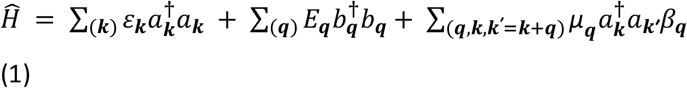

with energies ε_*k*_ = ℏω_*k*_,*E*_*q*_ = ℏ*Ω*_*q*_ (ℏ = Planck’s constant; ω, *Ω* = angular frequencies) depending on wavevectors *k, q*. A theorem states that the dynamics of *a*_*k*_, *b*_*q*_ can be kept separate from external energy–pump operators when a second-order coupling is retained^50^. The interaction energy, μ_*q*_ = χ sin^2^(**q** ⋅ **a/**2) Λ_*q*_, includes the unit cell vector **a**, the length scale Λ_*q*_ ≡ (ℏ**/**2*MΩ*_*q*_)^1**/**2^ and the lattice displacement operator 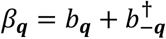. The mass scale *M* specifies the total number of atoms participating in the process, the anharmonic force χ quantifies how much the local excitation energy vary upon deforming the whole protein structure^44^. Now consider a correlated movement in the ground state (*q* = **g**) at a frequency *Ω*_**g**_. It is regular practice (SI–4c) to deduce the laws of motion in Heisenberg’s representation, getting an oscillatory equation for the average displacement *D*_**g**_ ≡ ⟨β_**g**_⟩ driven by the coupling interaction:

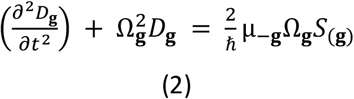

with 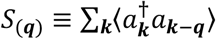 and *t* being the time. The absolute value of mean atomic displacements in this oscillatory steady–state is given by 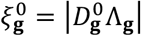, i.e. the stationary component of *D*_g_ times the characteristic length. The final result (SI–4c):

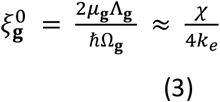

comes from replacing the acoustic dispersion law in *Ω*_**g**_,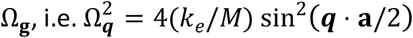, where *k*_*e*_ is a lattice spring constant. Eq. (3) is a classical relation: *atomic displacements arise from a balance between the harmonic force* (represented by *k*_*e*_) *experienced by atoms within the lattice and the anharmonic force* (χ) *that mediates the coupling between the lattice and local excitations* (Figure S5b). Using typical orders of magnitude in Davydov’s theory^48,51^, *k*_*e*_ = (10 − 20) N/m (H–bond) and χ = (10 − 80) pN, we get 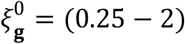 pm and the representative magnitude 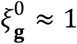 pm, in agreement with the observation. Optical effects should contribute to shifts ≤ 0.1 pm (SI–4d) and are regarded here as secondary. In summary, Eq. (2), resolved by a cosine wave (SI-4c), illustrates the overall inclination of the system, or its atomic constituents, to exhibit coherent oscillations. Consequently, coupled subsystems with (slightly or moderately) disparate oscillation frequencies may be anticipated to undergo synchronization processes, both in terms of Huygens synchronization or mode-locking scenarios.

## Predicted atomic shifts are more realistic in the presence of entrainment processes

We can show that the agreement with experiments improves when both large–scale and local excitations further entrain their motion. It suffices to include Heisenberg’s equations for the displacement of the local quasiparticle (SI–4e), with operator 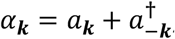, characteristic length λ_*k*_ = (ℏ**/**2*m*ω_*k*_)^1**/**2^, a mass scale *m* for the involved amino acid residues, and a reasonably constant dispersion law, ω_*k*_ ≈ ω_*a*_ ≡ ε_*a*_**/**ℏ ^52,53^. We attain three steady states (SI–4f): one corresponds to the former displacement, 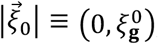; the other two involve a single–particle and a lattice component, 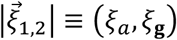, which satisfy 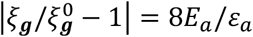, where *E*_a_ = *m***/**2 (ω_*a*_ ξ_*a*_)^2^ is the potential energy of a harmonic oscillator shifted by ξ_*a*_. Two complementary out–of–equilibrium regimes emerge: one phononic, with larger displacements than in classical states 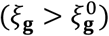; the other of a more local nature, with an opposite trend 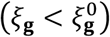. They widen the previously deduced range (i.e. 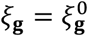). Employing ω_*a*_ ∼1 THz, *Ω*_**g**_ = (0.1 − 1) THz, χ = 63 pN, *k*_*e*_ = 13 N/m^48,53-56^, yields ξ_*a*_ ≈ (0.1 − 17) pm and ξ_**g**_ ≈ (1 − 13) pm for *M* ≈ (10^4^ − 10^5^) amu, and *m* ≈ (10^2^ − 10^6^) amu. When *m* = (57.05 − 186.2) amu (Gly-Trp) and *M* ≈ 10^5^ amu, we obtain in particular that ξ_*a*_ ≈ (10 − 20) pm, and ξ_**g**_ ≈ 1.5 pm.

Refining Eq. (3) to accommodate atom displacements ranging from fractions of pm to a dozen pm is an excellent result. In fact, when detected in different protein regions, their magnitude is elevated by ∼ (2 − 10) pm in comparison with control values (Figure 3g). This aligns with our steady–state predictions in the phonon deformation regime 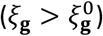 and with the emerging correlations between atomic shifts.

## Atom fluctuations (B-factors) in a squeezing process

The experimental behaviours of atomic fluctuations appear to be markedly irregular. A description of their collective trend can be provided by the frequency variation experienced by local excitations upon coupling with phonons, leading to an energy redistribution along the protein chain (SI–4g). Accordingly, the average B-factor, instead of the expected decrease produced by a higher order (i.e. larger correlations), may even increase. A second aspect can arise from the phonon kinetics (*b*_*q*_). In transitioning from thermal number to pure coherent states, quasiparticles will traverse through antagonistic configurations at intermediate molecular order, appropriately identified here as coherent squeezed states^57,58^. Squeezing is a purely quantum effect: the uncertainty in one quantity (e.g. *q*) lowers at the expense of raising the uncertainty in another (e.g. its conjugate momentum), and can play an important role in entrainment phenomena^59^.

To let this process emerge, we study the dynamics of 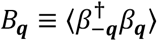 upon a lattice shift 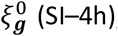, getting an oscillatory law ruled by the 2–mode quadrature squeezing operator, 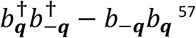. The final expression of the *B*-factor shows an oscillatory part which evolves as a decaying periodic function (SI–4i, SI–4l). Its amplitude has two main orders of magnitude (classical, c; semiclassical/quantum, sq):

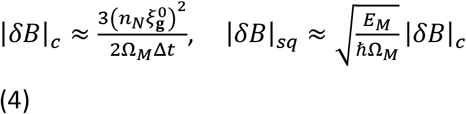

that depend on the number of atoms (*n*_*N*_) and the time scale (Δ*t*) involved in the process^60^, while Ω_*M*_ is a (Debye-like) cut-off frequency and Λ_*M*_ is the typical length scale in the coupling energy related to Ω_M_ (Eq. 1, last term). The energy 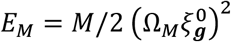 depicts a harmonic oscillation involving *n*_*N*_ atoms, of total mass *M*, shifted by 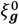. Eq. (4) exhibits an increase with both increasing *n*_*N*_ and the softness of the crystal system (1**/**Ω_*M*_), where the atomic displacements 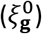 represent an additional fluctuation source. If Δ*t* = (1 fs − 1 ms), *n*_*N*_ ∼ (1 − 10^6^), 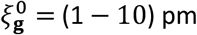 pm, and Ω_*M*_= 5 THz, then |δ*B*| turns out to range from a few cents to a few tens of Å^2^ (SI–4l), corresponding well with experimental magnitudes ∼ (10^−2^ − 10) Å^2^. These estimates are more phenomenological in nature rather than absolute, as they require further details on protein dynamics to be fully comprehensive. However, they suggest a novel framework for interpreting atomic fluctuations and shifts in ultrafast experiments (Figure S6).

In summary, ultrafast Terahertz (THz) pump X-ray diffraction probe experiments were conducted on bovine trypsin crystals. We devised a resampling strategy for quantifying the uncertainty of atom displacements and for evaluating atomic pair correlations. No evidence of unit cell deformations was found in protein crystals 50 ps after the exposure to high-field THz pulses. *A strong link appears between direction/magnitude of displacements and fine–grained atom types, and a weak link between atom displacements and locations* (Figure S7, Movie S1). Several cases where the magnitude under pumping exceeds the control steady state value were identified. An analysis of atom pairs in resampled datasets revealed a relatively strong correlation between distal atoms, which diminished if the measurement followed a THz pulse. A THz periodic electric field (**E**) in molecular dynamics (MD) simulations may partly explain the role of electrodynamics in developing correlated shifts between similarly charged atoms but, especially in low-field regimes, they more likely arise from coherence, mode-locking entrainment and self-oscillation^61^. Correlated patterns between distal atoms, widely distributed in a scaffold that does not deform elastically as a whole, prompt a need to reassess current protein models. Our theoretical description focuses on the time–coherence of elastic domains near or at a steady state. The model achieves correct magnitudes of atom displacements, which improve when large–scale (lattice deformation) and smaller–scale quasiparticles (local excitations) entrain together.

Analysis of the B-factors raises a complex picture, reminiscent of a (quantum) “squeezing” process. This view, clearly, does not model the THz pulse induced transient stage before a steady steady state is restored. While a time interval of 50 ps may not be enough for atomic displacements to entrain again, a steady state is expected to reform within the cycle time of experiments (500 ms).

Overall, our measurements, simulations and models strongly suggest that protein symmetry extends beyond solution structures and propose a connection to the coherent dynamics of atoms and functional groups, as detected in crystals of bovine trypsin. Concerning the biological function of these groups, there has been speculation about which properties of peptides are crucial in determining their interaction with promiscuous proteins such as survivin. In this example, it appears that the sequence does not provide substantially more deterministic power compared to composition^62,63^. However, for an effective model, the composition must include a detailed description of functional groups in the amino acids, rather than simply considering the number of hydrophobic and hydrophilic amino acids, net charge, and charge density. In this study, coherent dynamics were observed in a crystal structure through femtosecond X-ray diffraction snapshots. We expect that our theoretical arguments could be generalized to different scales of organization, ranging from interactions of proteins and peptides to the microscopic organization of multicellular organisms at a higher level. In this context, it is noteworthy that the embryonic scale structure of a complex organism like a dog is influenced by the number of repetitive functional groups in key proteins^64^. Future research is required to map other functional and structural implications of coherent protein dynamics in biological organization.

## Acknowledgements

Experiments were performed at the FemtoMAX beamline of the Max IV synchrotron. This work was supported by the Röntgen-Ångström Cluster Framework (2015-06099). S.A.M acknowledges financial supports from the Institute for advanced Neutron and X-ray Science (LINXS/LU) and the Croatian Science Foundation (HrZZ) through the project IP-2022-10-3456. The authors gratefully acknowledge Daan Frenkel from Cambridge University for reviewing the manuscript and highlighting the intriguing correlation between distal atoms and allostery. Computations were enabled by resources provided by the Swedish National Infrastructure for Computing (SNIC) at the PDC Center for High Performance Computing, KTH Royal Institute of Technology, partially funded by the Swedish Research Council through grant agreement no. 2018-05973. J.L. acknowledges support from the Swedish Research Council (VR, Grant No. 2015-06115). This project has received funding from the European Union’s Horizon 2020 research and innovation programme under grant agreement No. 964203 (Long-range electrodynamic INteractions between proteinS’ — LINkS).

## Data availability statement

The initial diffraction data images have been submitted to the Zenodo database (DOI: 10.5281/zenodo.8155329). Additionally, the resampled and summed versions of these images, along with resampled reduced structure factor amplitudes, and 200 refined structures obtained from the resampled data sets will be made available on the Zenodo database upon acceptance of the manuscript. Furthermore, a representative sample of the THz pumped and reference structure will be deposited at the PDB database upon acceptance of the manuscript. Other data that support the findings of this study are available upon reasonable request and by contacting the corresponding authors.

## Code availability statement

All original code will be deposited at the Zenodo and Github databases and will be publicly available as of the date of publication. These include the python scripts utilized for cbf diffraction image generation, incorporating libraries such as numpy, scipy, pandas, and fabio. These scripts were built upon earlier work32 and will be made publicly available on GitHub once the manuscript is accepted. Data reduction was conducted using XDS, XSCALE, and XDSCONV, while the CCP4 package, specifically REFMAC5, was employed for data analysis and refinement. To enable efficient parallel analysis, Python and shell wrapper scripts were implemented and will also be openly accessible upon manuscript acceptance. For plot visualization, we relied on Python libraries matplotlib and seaborn. Pymol, with the “modevector” plugin, facilitated the visualization of molecular structures and directions. Additionally, the model analysis scripts will be deposited on GitHub once the manuscript is accepted. For image and video editing, as well as labelling, Inkscape and Openshot were utilized. The watermark will be removed from the published supporting media file.

## Inclusion and Ethics

We support inclusive, diverse, and equitable conduct of research.

