## Supplemental information for "Femtosecond X-ray snapshots reveal correlated displacements of specific distal atoms in a protein crystal"

### 1. Supplementary discussion

#### *a.) Solvent composition of trypsin crystals*

Starting with the initial model structure, the solvent content is determined to be 46%. Assuming the solvent is pure water, this corresponds to approximately 3360 water molecules in the unit cell, calculated as $0.46 \times N_A \times 55$ M × 55.1 Å × 58.9 Å × 67.9 Å, where $N_A$ represents Avogadro's constant. With the primitive orthorhombic unit cell containing four asymmetric units, each asymmetric unit is associated with roughly 840 water molecules. Within the asymmetric unit, there are 184 modelled ordered water molecules per protein. To ensure numerical stability during the refinement of various resampled data sets, weak electron density peaks have not been modeled as potential water molecules, even when the hydrogen bonding geometry to the peak is suitable for a water molecule. Therefore, the count of 184 molecules should be regarded as a conservative estimate for the observable ordered water molecules.

The remaining solvent region, referred to as the "bulk solvent" in crystallography, encompasses the rest of the water molecules, approximately 656 in number. This region differs from the aqueous bulk in solution studies, due to the presence of the crystal lattice. In crystallographic investigations, the bulk solvent region is often approximated with uniform electron density, impeding a detailed analysis of this masked volume. The total number of water molecules can be compared to the 1629 non-hydrogen protein atoms, taking into account alternative locations counted only once.

#### *b.) Expected temperature change upon THz pulse absorption*

Assuming the crystal shares similar density and heat capacity characteristics with water, a full absorption and thermalization of a 4 µJ THz pulse would result in a peak temperature increase of $\frac{4\mu J}{4184\,\frac{J}{kgK}\times(300\mu m)^3\times 1\frac{kg}{L}}$ = 35 mK. Considering that not all energy is absorbed in the crystal, thermalization requires time and dissipation starts immediately, this 35 mK temperature increase also represents the maximum achievable change. At most, this temperature rise would alter the conformational state population by 2.4%. This estimation is derived by correlating the temperature rise with a change in equilibrium constant of the conformational state population using the van't Hoff equation, considering enthalpy.

Examining the impact of a 35 mK temperature increase on the most extensively studied conformational change in proteins, namely thermal unfolding, provides further insight. Proteins the size of trypsin typically exhibit an unfolding enthalpy change of around 1000 kJ/mol[1]. The phase in the unfolding process most sensitive to temperature changes lies near the melting temperature, where the folded and unfolded state populations are equal. Even in the unlikely scenario of substantial enthalpy change and the system being precisely at the melting temperature, a 35 mK temperature rise would merely shift the conformational state population by 2.4%.

$$K_1 = \frac{[U]}{[F]} = 1$$

$$K_2 = K_1 \times \exp\left(\frac{1}{R} \times 1000\,\frac{kJ}{mol} \times \left(\frac{1}{300.000K} - \frac{1}{300.035K}\right)\right) = 1.05$$

$$K_2 = \frac{1+\alpha}{1-\alpha}$$

$$\alpha = 0.024$$

It is also unrealistic to anticipate a conformational change occurring at the same speed as thermalization. Therefore, achieving the new conformational equilibrium would require orders of magnitude more time than the 50 ps delay time allotted for the protein system. From a purely equilibrium thermodynamic perspective, at this THz pulse energy level, the induced temperature variation is insufficient to noticeably influence even an extremely sensitive conformational equilibrium.

#### 2. Experimental Methods

##### *c.) Protein Crystallization*

Bovine trypsin was dissolved in 30 mM HEPES at pH 7.0, 3 mM $CaCl_2$ and 6 mg/ml benzamidine hydrochloride hydrate to a concentration of 60 mg/ml. Crystals were grown using the hanging drop vapor diffusion method at room temperature. The crystallization drop was prepared by mixing 5 µl protein solution and 5 µl of precipitant solution (18% PEG8000, 50 mM HEPES at pH 7.0, 0.2 M ammonium sulfate, 3 mM $CaCl_2$ and 6 mg/ml benzamidine hydrochloride hydrate). Crystals of the approximate size of 300 µm were harvested after three days by polymer loops (MiTeGen), and stabilized at room temperature by a plastic capillary (MiTeGen) [2]. All chemicals and the protein were purchased from MilliporeSigma.

##### *d.) THz pulse generation and synchronization*

THz pulses were generated by optical rectification in 4-N,N-dimethylamino-4'-N'-methyl stilbazolium tosylate (DAST) crystals (Swiss Terahertz)[3]. A 800 nm, 12 mJ, 45 fs Ti:sapphire laser (Red Wyvern from KM Laboratories) was adopted for pumping an Optical Parametric Amplifier (TOPAS HE from Light Conversion) to generate approximately 1 mJ, 1500 nm pulses at 100 Hz repetition rate. Subsequently, 0.6 mJ of the laser light excited the DAST crystals, which in turn produced 4 μJ, 2.1 THz radiation (with FWHM of 2 THz) after filtering residual infrared (IR) radiation (Swiss Terahertz) and focused by a TPX lens (50 mm focal length and 50 % transmission, Tydex) to an approximately 500 μm spot size at the protein crystal (full width at half maximum). The THz field was linearly polarized parallel to the table top and the magnetic field was parallel to the crystal spindle axis. Schematics of the beamline is displayed in Figure 1a and 1b in the main text. The electric field strength over the protein crystal was estimated to be about $1.2 \times 10^2$ V/μm. THz field reflections at the crystal surface were not considered in this estimation.

Both the IR pump laser and electron gun were synchronized to a common 3 GHz radio frequency (RF) signal with an approximate root mean square (RMS) accuracy of 50 fs. However, the timing was limited by a pulse-to-pulse jitter of typically < 1 ps[4]. The temporal overlap was established by a Hamamatsu G4176-03 photodiode placed behind the sample and the timing difference was measured between the direct X-ray pulse and scattering of the pump laser pulse from the sample holder loop in the centered position.

The THz pump pulse had a 100 Hz repetition rate and each 50$^{th}$ pulse (incoming at every 0.5 s) was nearly coincidental with the X-ray probe pulse, operating at a 2 Hz repetition rate. A millisecond (ms) shutter was synchronized with the Pilatus detector. A DG645 pulse (delay) generator (Stanford Research Systems) selected every second a synchronized pump pulse leading to the odd frames as non-irradiated (diffraction images $DI_{off,i}$) and to even frames as irradiated ($DI_{THz,i}$). To achieve the desired pattern, a pulse was triggered on the raising edge of the EN OUT signal of the Pilatus detector and started the ms shutter opening pulse (duration 100 ms). This allowed the ms shutter to open in advance of the next pump pulse. A hold-off period of 0.75 s ignored the next Pilatus trigger. This also meant that the first and every odd frames were recorded with the ms shutter closed ($THz_{off}$). For those frames which were irradiated, the pump-probe scheme consisted of a single-cycle, broadband THz pulse followed by a single 150 fs X-ray pulse, and separated by a time delay of 50 ± 1.3 ps. A schematic illustration of the timing scheme is included in Figure 1c.

*e.) X-ray diffraction data collection strategy at the FemtoMAX beamline*

The X-ray diffraction data collection strategy has been described elsewhere[2], where diffraction from a non-THz pumped crystal was determined. The crystal was studied by a total angular range of 134.4°. We measured 100 X-ray diffraction images per rotation position, while the crystal was kept still (0.1° separation between each position). Each image corresponded to measurements from an individual X-ray pulse. The second and every even numbered ones were recorded by a THz pump pulse preceding the probe X-ray pulse ($DI_{THz,i}$). Odd numbered diffraction images were recorded without a preceding THz pulse ($DI_{off,i}$).

The X-ray beam was lost due to regular storage ring refills and irregular problems, but the detector kept recording to fill the blocks of 100 images per position. Thus, there were empty images afterwards. Those non-empty were identified by a custom-made Python script, using numpy and pandas packages[5,6] in which images with an average pixel count above a certain threshold (i.e. 0.005 counts/pixel) were selected. In rare cases entire rotation positions were compromised and no images were generated. The empty ones were removed in pairs whenever some issue was identified. For example, if an even numbered image was missing then the following odd one was removed as well even though it contained useful data (and vice versa). We finally obtained an equal number of (as consecutive as possible pairs of) odd and even images in datasets.

Ideally, 50 odd and 50 even numbered images were produced per crystal rotation. For testing the robustness of data quality indicators determined from exported data, and describe a distribution rather than point estimates, we resorted to a resampling strategy. To avoid the need for replacements without large loss of data quality, only half of the DI was used for generating a resampled dataset. Odd and even numbered images were resampled in pairs. For instance, if image #3 was assigned by chance to odd dataset #1, then the even dataset #1 automatically acquired image #4. A strict pairing of odd and even images was mandatory to reduce systematic errors that might distort the difference attributed to THz irradiation. The equivalent pixel values were summed over each individual set and exported as .cbf files by the FabIO package[7]. Resampling was performed without replacement, one image was used maximally once in sums of diffraction images such as $\sum_i^j DI_{k,i}$ (k = THz, off). On the other hand, since summation of identical images disrupts the counting statistics in the data processing software, resampling with replacement was disallowed.

Note that, at positions with less than 50 images per crystal rotation, resampling was performed over a smaller pool of original images. Since approximately 25 % of images are shared among resampled datasets, the resampled sets were not statistically independent of each other within the odd and even groups, i.e. for any odd-odd and even-even pairing. Handling of data is illustrated in Figure 1d.

*f.) Data reduction and refinement*

A Python script performed an automated processing of individual datasets, resorting to a combination of the following software: X-ray Detector Software (XDS), XSCALE, XDSCONV and REFMAC5 [8-10]. Sums of diffraction images $\sum_i^j DI_{THz,i}$ and $\sum_i^j DI_{off,i}$ were indexed and integrated in XDS. Default settings for non-anomalous diffraction were used in the subsequent scaling by XSCALE, merging and conversion operations from intensities to structure factor amplitudes were performed by XDSCONV. In the final XSCALE step, a fraction of resampled data sets exhibited deviating statistics, particularly noticeable from the 4.55-3.72 Å resolution bin. Here, a discontinuous jump of >20% in $R_{merge}$ occurred, instead of the typical smooth increase to less than 10% seen in most data sets. When a data set displayed deviating statistics, both the THz and off data sets were removed with the corresponding resampling ID. This scaling artifact was not strictly associated with either the $THz_{on}$ or $THz_{off}$ data sets. After removing these data sets (resampling IDs: 8, 10, 16, 17, 19, 22, 27, 29, 31, 32, 35, 41, 43, 53, 64, 67, 68, 75, 76, 78, 79, 87, 93, 96, 98 and 99), 74 $THz_{on}$/$THz_{off}$ data set pairs remained.

An amount of 74 $F_{THz,n}$ and 74 $F_{off,n}$ data were independently refined by means of the same starting model and REFMAC5 software. First, a maximum-likelihood rigid body refinement was conducted over 100 cycles, followed by 100 cycles of a maximum-likelihood restrained isotropic refinement. Refinements operated with no prior phase information and automatic weights between structure factors and geometric restraints. The resulting crystallographic models represent point estimates of atomic positions and individual isotropic atomic B-factors as part of 74 $M_{THz,n}$ and 74 $M_{off,n}$ models. The error bar stands for the standard deviation of point estimates. Crystallographic models were visualized by Coot[11]. Isotropic B-factor is defined as $B_{iso} = 8\pi^2\langle u^2\rangle$[12], where $\langle u^2\rangle$ is the mean atomic displacement expressed in $Å^2$. We summarized the distributions of data quality indicators, produced from the scaling and refinement in Table S1, by carrying out arithmetic means and standard deviations.

*g.) Model analysis*

Resampling was performed over pairs of images $DI_{THz,i}$ and $DI_{off,i}$, which ultimately resulted in the crystallographic models $M_{THz,n}$ and $M_{off,m}$. In this notation, symbols $M_{a,THz,n}$, $M_{b,THz,n}$ and $M_{a,off,n}$, $M_{b,off,n}$ are used to compare pairs of atoms, *a* and *b*, picked up from $M_{THz,n}$ and $M_{off,n}$. Vector deformation fields $\boldsymbol{M}_{THz,n} - \boldsymbol{M}_{off,m}$ were thereafter calculated for the displacement of mean positions upon $n = m$. Control fields $\boldsymbol{M}_{off,n} - \boldsymbol{M}_{off,m}$ were generated from $M_{off,n}$ and $M_{off,m}$ with $n = m + 50$ and $1 \leq m \leq 50$ ($n, m$ natural numbers). A similar point can be made for the B-factor changes, with scalar deformation fields $B_{THz,n}$ - $B_{off,n}$.

By using IOTBX of the CCTBX package[13], we replaced the B-factors values in one of the refined protein models with B-factor differences, and coloured the protein model in Pymol[14] according to the obtained numerical value. In addition individual atom coordinates were substituted by new coordinates, calculated from scaling the average displacements 100 times. The Pymol script "Modevectors"[15] then was implemented to visualize vectors in figures.

By the seaborn package[16] in Python, we also benefitted from a non-supervised hierarchical clustering method to find connections between mean atomic displacements. We first reduced our data by calculating the mean deformation field $\langle \boldsymbol{M}_{THz,n} - \boldsymbol{M}_{off,n} \rangle$ on each individual atom. Clustering then was performed through the cityblock distances ($\ell_1$ norms, built upon absolute values) between the mean deformation fields, i.e.:

$$d(a,b) = \left\| \langle \boldsymbol{M}_{a,THz,n} - \boldsymbol{M}_{a,off,n} \rangle - \langle \boldsymbol{M}_{b,THz,n} - \boldsymbol{M}_{b,off,n} \rangle \right\|_1 \qquad \text{(S1)}$$

where *a* and *b* are the two atoms to be confronted. The average linkage clustering method was used to perform the calculation.

#### 3. Computational Methods

##### *a.) Survey of human homooligomeric proteins*

The survey was conducted using the rcsbsearchapi python library (rcsbsearchapi.readthedocs.io) [17]. The initial selections always utilized the following query attributes: rcsb_struct_symmetry.kind == "Global Symmetry", rcsb_entity_source_organism.ncbi_scientific_name == "Homo sapiens", and exptl.method == "X-RAY DIFFRACTION". Additionally, the attribute rcsb_struct_symmetry.symbol was used to select one of the point groups analyzed in this study. This was done in conjunction with the attribute rcsb_entry_info.deposited_polymer_entity_instance_count, where the count was restricted to either 1 (alternative 1) or integer multiples of the minimum chains corresponding to the selected symmetry (C2: 2n, C3: 3n, C4: 4n, D2: 4n, D3: 6n, D4: 8n, alternative 2). Subsequently, the uniprot entry of every chain was gathered from each returned PDB entry. If a retrieved PDB entry contained more than one UniProt entry or did not contain the exact number of requested chains, it was discarded. In the final summary, each unique uniprot entry was counted only once.

##### *b.) Molecular dynamics (MD) simulation in the presence of a THz electric field*

MD simulations were performed by the Gromacs simulation package[18-20]. The same initial position was used in all simulations, as the difference in resampled positions (coordinates) is smaller than random initial condition differences in MD. Ten configurations were initially generated by rotating the starting coordinates randomly by choosing three angles between 0° and 360°, and generating a box of (7.624 x 7.624 x 7.624) $nm^3$ around the rotated models.

Models were solvated with pre-equilibrated water and the ion strength was set to the equivalent of 150 mM NaCl in addition to neutralization of the net protein charge with $Cl^-$ ions[21]. The CHARMM36[22-24] force field was used in all calculations. To be able to simulate the protein realistically by this force field, it was necessary to remove the non-protein residues SO4 and BEN.

Prior to carrying out the simulations, 5000 steps of energy minimisation were run, employing the Steepest Descents algorithm. This yielded a maximum force of $1.46 \times 10^3$ kJ $mol^{-1}$ $nm^{-1}$. Following the minimisation, equilibration was conducted as follows. First, a short (20 ps) run was made while keeping the non-hydrogen protein atoms restrained with a force constant of $10^3$ kJ $mol^{-1}$ $nm^{-2}$. Thereafter, a 5 ns equilibration simulation was run unconstrained in each system.

During equilibration, temperature was kept constant (T = 293 K) by means of the velocity-rescaling algorithm ($\tau_T$ = 0.1 ps)[25]. Pressure was coupled to an external bath by means of Berendsen's coupling algorithm[26] ($P_{ref}$ = 1 bar, $\tau_P$ = 1 ps) during equilibration, and with the Parrinello–Rahman algorithm[27] afterwards. van der Waals forces were truncated at 1.0 nm with a plain cut-off. Long-range electrostatic forces were treated by the particle mesh Ewald's summation method[28]. Dispersion correction was applied to energy and pressure. The Cα RMSD was small (< 0.15 nm) in all cases.

Following equilibration, six sets (6 × 10) of simulations were carried out: one with no electric field and the others by using non-pulsed field of 35, 105, 175, 250, and 350 MV/m with frequency 2 THz. Each run's duration was 10 ns, during which snapshot structures were saved every 10 ps and systems were analysed to view any deformations; None such were observed. To examine correlation between atomic motions, runs were extended by 20 ps (100 times the period of the field), and every frame was saved. The covariance matrix (cov) was generated by the utility gmxcovar with the option -xpma. Thus, the atomic covariance matrix contained, for each atom pair, the sum of Cartesian xx, yy and zz covariances. A correlation matrix (corr) was finally inferred from:

$$corr(X_i, X_j) = \frac{cov(X_i,X_j)}{\sqrt{cov(X_i,X_i)\cdot cov(X_j,X_j)}} \quad \text{(S2)}$$

*c.) Notes on comparing covariance matrices, obtained from Molecular Dynamics (MD) simulations, with resampled experimental data.*

MD simulations predict atom trajectories of one or limited numbers of protein molecules. Covariance and correlation can be calculated between the time dependent coordinates of atom pairs. Due to the significance of covariance, the order of time points in the trajectory is not important for defining the "cov" matrix and any permutation of structures yield the same results. Similarly, if we only consider randomly sampled subsets of the trajectory, they can provide an estimate of covariance of the entire evolution. When a large number of proteins follow independent trajectories in the crystal lattice, correlation should drastically reduce and the contribution expected to remain is coming from non-independent atom displacements. If we experimentally record nearly instantaneous snapshots of a random or deterministic process in arbitrary order along some reaction or vibrational coordinate, in principle we should be able to capture a similar information, and the simulated and experimental atomic covariances can be confronted. Resampled data sets thus could represent slightly different positions along a process coordinate.

In crystallographic refinements, hydrogen (H–) bonds are not directly restrained, short distances between hydrogen bond acceptors and donors are simply tolerated. Restraints are confined to covalent bond lengths, angles, and in specific cases to torsion and improper angles, preventing steric clashes assimilated to the repulsive part of a Lennard-Jones potential for non-covalent interactions. Moreover, at high resolutions and well-ordered regions, such restraints are downweighted relative to the likelihood based on the amplitudes of crystallographic structure factors. Positions of distal atoms can be regarded independent of restraint effects.

### 4. Theoretical Methods

*a.) Dynamic description of Fröhlich's system*

In his seminal paper[29], Fröhlich introduced the energy dissipation rates from 2–body ($L_i$) and 3–body ($\mathbf{L}_{ik}$) interactions of a biological system with the surrounding bath states:

$$\frac{dn_i}{dt} = s_i + \phi_i L_i - \sum_k \lambda_{ik} \mathbf{L}_{ik} \quad \text{(S3)}$$

where, for each vibration mode $i$, $n_i$ is the number of quanta and $s_i$ is the energy supply rate (here, indices $i$ or $k$ are not wavenumbers or wavevectors). The terms $\phi_i(T)$ and $\lambda_{ik}(T)$ are the linear and nonlinear temperature–dependent coupling rates between heat bath and one or two system states, respectively. The former is reminiscent of a linear damping term in Langevin's approach, and is meant to quantify dissipation effects. The latter governs energy redistribution among modes. The mathematical forms of $\mathbf{L}_{ik}(n_i, T)$ and $L_i(n_i, T)$ were chosen so as to return a Bose-Einstein distribution in the absence of external pump energy, $\mathbf{L}_{ik}(\bar{n}_i, T) = 0$ and $L_i(\bar{n}_i, T) = 0$:

$$L_i = \frac{n_i}{\bar{n}_i} - 1 \quad \text{(S4)}$$

$$\mathbf{L}_{ik} = (p_{k-i} - 1)n_i n_k + p_{k-i}(n_i - n_k) \quad \text{(S5)}$$

in which $\bar{n}_i = (e^{\beta\hbar\omega_i} - 1)^{-1}$ is Planck's distribution function for the mean phonon occupation number, $\beta = 1/(k_B T)$ is the reciprocal of Boltzmann's energy, $\hbar$ is Planck's constant and $p_{k-i} \equiv e^{-\beta\hbar(\omega_k - \omega_i)} = \bar{n}_{k,i}/(1 + \bar{n}_{k,i})$. Interestingly, on interpreting $p_{k-i}$ as a Boltzmann's probability, the expression for $-\mathbf{L}_{ik}$ implies for $n_k = 1$ a rate contribution of the Markov-type[30], with a classical transition probability from state $i$ to state $k$. This hint was looked through in a former work[31], proving mathematically that Fröhlich's view basically leads to a master equation which implicitly assumes a time–homogeneous Markov process. Generally, the evolution of polar mode populations in Eqs. (S3–S5) may be derived satisfactorily from the nonequilibrium statistical operator method in a quantum relaxation theory at second order[32]. On such basis, Fröhlich inferred a coherent collective state for a nonequilibrium open system, where one or a few modes get overpopulated at the expense of the other vibrational levels. In a nutshell he had found that, on increasing the power input supply, the energy of longitudinal electric vibrations is channelled into a non-thermal condensate, similar to Bose-Einstein's. It should be pointed out that, in principle, the identification of the energy source may be arbitrary, as it depends on the mutual definition of system and environment. We won't cope in this paragraph with this aspect, nor with a discussion of chemical potentials (crucial to establish the onset of condensation), but enter Fröhlich's regime and recast it mathematically in the simple case of three dipolar modes, one of which representing the ground state. Any extension to a larger ensemble of modes will follow an identical conceptual route. For formal convenience, we start from Eq. (S3), rewritten as:

$$\frac{\mathrm{d}n_i}{\mathrm{dt}} = \bar{\sigma}_\mathrm{i} - \phi_\mathrm{i} n_i - \sum_{k\neq i} \Lambda_\mathrm{ik}\,[n_i n_k + (1 + \bar{n}_{i,k})n_i + \bar{n}_{i,k} n_k] \tag{S6}$$

with $\bar{\sigma}_\mathrm{i} \equiv s_i + \phi_\mathrm{i}\bar{\mathrm{n}}_\mathrm{i}$, $\Lambda_\mathrm{ik} \equiv p_{i-k}\lambda_{ik}$, and develop Eq. (S6) upon:

$$\bar{n}_{i-k} + \bar{n}_{k-i} + 1 = 0,\ \Lambda_{\mathrm{i}<k} \equiv -\lambda_{\mathrm{i}<k},\ \Lambda_{\mathrm{i}>k} \equiv \lambda_{\mathrm{i}>k} \tag{S7}$$

where the first relation is a basic mathematical property, and the last two come from the definition of $\Lambda_\mathrm{ik}$. In this way, the final equation system only involves coefficients $\lambda_{ik} > 0$. For high enough temperatures (say, $T$ > Debye's Θ), one may work for simplicity with $\bar{n}_{i,k} \approx k_B T/\hbar(\omega_k - \omega_i) \approx -\bar{n}_{i,k}$.

Accordingly, when three states $|0\rangle$, $|1\rangle$, $|2\rangle$ are regarded with $\omega_0 < \omega_1 < \omega_2$, Fröhlich's condensation would be described by:

$$\frac{dn_0}{dt} = \bar{\sigma}_0 + \left(\lambda_{01}\bar{n}_{0,1} + \lambda_{01}\,\bar{n}_{0,2} - \phi_0\right)n_0 + \lambda_{01}n_0 n_1 + \lambda_{02}n_0 n_2$$

$$\frac{dn_1}{dt} = \bar{\sigma}_1 - \lambda_{01}\bar{n}_{0,1}n_0 - \lambda_{01}n_0 n_1 + \lambda_{12}n_1 n_2 \tag{S8}$$

$$\frac{dn_2}{dt} = \bar{\sigma}_2 - \lambda_{02}\bar{n}_{0,2}n_0 - \lambda_{02}n_0 n_2 - \lambda_{12}n_1 n_2$$

As the ground state $|0\rangle$ is expected to become more and more occupied, terms in $n_0$ and/or $n_\mathrm{i}n_\mathrm{k}$ were only retained along with energy supply rates $\sigma_k$. These model equations, despite being essential, captures the main phenomenological features. While there certainly exists a range of coefficient values such that $n_2 < n_1 < n_0$, distribution of signs in both linear and nonlinear terms actually implies that the phonon number in $|0\rangle$ is detrimental to $|1\rangle$ and $|2\rangle$. Under steady conditions (suffixed by *), the number of quanta in the ground state becomes:

$$\phi_0 n_0^* = \sum_{k\geq 0}\sigma_k \equiv \Sigma \tag{S9}$$

or, carrying out every term:

$$n_0^* = \bar{n}_0 + \frac{1}{\phi_0}\sum_{r>0}\phi_r\bar{n}_\mathrm{r} + \frac{1}{\phi_0}\sum_{k\geq 0} s_k \tag{S10}$$

Consistently with the original work[29], this relation expresses that $|0\rangle$'s phonon number linearly increase with the (total) supply rate. It equals the equilibrium value augmented by a weighted sum of corresponding values in higher energy levels. Furthermore, $n_0^*$ turns out to be enhanced by lowering the damping coefficient of the $|0\rangle$ channel.

In the linear stability analysis a full solution can be given in explicit form once we notice that, eventually, $n_k n_h \ll n_i n_q$, with $k, h, q \neq i$. The steady state solution $\mathbf{n}^*$ then has the following components:

$$n_0^* = \frac{\Sigma}{\phi_0}, \qquad n_k^* = \frac{\phi_0 s_k}{\lambda_{0k}\Sigma} - \bar{n}_{0k} > 0, \quad (k = 1,2) \tag{S11}$$

the addition of vibrational modes with larger $k$ descending from alike recursive rules. Eqs. (S8) can be recast in terms of the vector notation $\mathbf{n} = (n_0, n_1, \dots)$ and linearized upon $\mathbf{n} = \mathbf{n}^* + \delta\mathbf{n}$:

$$\left(\frac{d\delta\mathbf{n}}{d\tau}\right) = \mathbf{U}^*\delta\mathbf{n} + \mathcal{O}(\delta\mathbf{n}^2) \tag{S12}$$

where $t/\tau \equiv \vartheta$ denotes some characteristic time, $\mathbf{U}^* = \mathbf{U}(\mathbf{n}^*)$, and:

$$\mathrm{U}_{qh}(\mathbf{n}) = \frac{\partial}{\partial \mathrm{n}_h}\left(\frac{d\mathrm{n}_q}{d\tau}\right) \tag{S13}$$

The final result is quite simple and function of the phonon number in the condensate as:

$$\boldsymbol{U}^* = \vartheta n_0^* \begin{pmatrix} s_1/{n_0^*}^2 & -\lambda_{01} & -\lambda_{01} \\ s_2/{n_0^*}^2 & \lambda_{01} & 0 \\ s_3/{n_0^*}^2 & 0 & \lambda_{02} \end{pmatrix} \tag{S14}$$

with characteristic eigenvalue equation:

$$\|\mathbf{U}^* - \alpha\mathbf{I}\| = 0 \tag{S15}$$

that reads:

$$\alpha^3 + \vartheta^3[\sigma_0\,\phi_0^2 + (\lambda_{01} + \lambda_{02})\Sigma^2]\,\alpha^2 + \vartheta^6\Sigma^2\phi_0^2\left[\lambda_{01}(\sigma_0 + \sigma_1) + \lambda_{02}\,(\sigma_0 + \sigma_2) + \lambda_{01}\lambda_{02}\left(\frac{\Sigma}{\phi_0}\right)^2\right]\alpha = -\vartheta^9\Sigma^5\phi_0^2\lambda_{01}\lambda_{02} \tag{S16}$$

Eq. (S16) defines a tough problem in a $\mathfrak{R}^{11}$ parametric space, whose full discussion is unimportant here. However, the real part of each eigenvalue is $\mathfrak{R}\{\alpha_i\} \leq 0$, which means that the condensate is stable or asymptotically stable. Now that condensation growth is framed into a formally consistent model, we can resume the original Fröhlich’s work and resort to a common energy supply rate per mode ($s$), damping ($\phi$) and redistribution terms. This allows to work with the natural time scale $\vartheta \approx 1/s$ and to recast Eq. (S16) into:

$$\alpha^3 + [\phi_s^2(1 + \phi_s^2\bar{n}_0) + 2l\phi_s(3 + \phi_s\bar{n})^2]\,\alpha^2 + \left\{l^2\phi_s^2(3 + \phi_s\bar{n})^3\left[1 + \left(\frac{\phi_s}{l}\right)\left(4 + \phi_s(\bar{n}_0 + \bar{n})\right)\right]\right\}\alpha = -\,\phi_s^4 l^2(3 + \phi_s\bar{n})^5 \tag{S17}$$

where $\bar{n} = \sum_k \bar{n}_k$ and $\phi_s \equiv \phi/s$, $l = \lambda/\phi$ are two dimensionless constants. Since atoms and residues in our system turn out to experience a sizable energy redistribution, we develop the linear stability results in the limit $\phi_s \ll 1$, $l \to \phi_s$, compatible with the formation of 'strong condensates'[33] and consider the polynomial coefficients in Eq. (S17) expanded to the lowest orders in $\phi_s$:

| | |
|---|---|
| $1$ | $(\alpha^3)$ |
| $3^2 l\phi_s + \mathcal{O}(2)$ | $(\alpha^2)$ |
| $3^3 l^2\phi_s^2 + \mathcal{O}(3)$ | $(\alpha^1)$ |
| $3^5 l^2\phi_s^4 + \mathcal{O}(5)$ | $(\alpha^0)$ |

An analysis of the roots of Eq. (S17) endowed with the above coefficients returns the three eigenvalues:

$$\alpha_0 = 3l\phi_s\left(\sqrt[3]{1-\frac{9\phi_s}{l}}-1\right) < 0 \quad \text{(S18)}$$

$$\alpha_{1,2} = -\frac{3}{2}l\phi_s\left(\sqrt[3]{1-\frac{9\phi_s}{l}}+2 \pm i\sqrt{3}\sqrt[3]{1-\frac{9\phi_s}{l}}\right) \quad \text{(S19)}$$

which, for $l \to \phi_s$, reduce to one real negative and two purely imaginary eigenvalues:

$$\alpha_0 \approx -9l\phi_s\,, \quad \alpha_{1,2} \approx \mp i\frac{1}{\sqrt{3}}\alpha_0 \quad \text{(S20)}$$

To sum up, the long–time phenomenological behaviour captured by this Fröhlich's condensation model, upon high enough supply rates and redistribution between vibrational modes, tends to be oscillatory near/at a marginally stable steady state. Values of frequency and time constant are clearly heuristic, as their direct link with quantities in real protein experiments is hardly achievable.

*b.) Near-threshold thermodynamic interpretation*

Fröhlich's condensation is an out-of-equilibrium phenomenon, whose energy threshold is tough to be predicted in advance for a real system such as a protein crystal. However, one may always imagine to work nearby the onset of marginal stability, and attempt a near–equilibrium description. Temperature changes upon application of the THz field are unimportant in our experiments, while volume variations ($\Delta V$) of the unit cell turn out to be smaller than values which can be resolved by X-ray crystallographic tools[34]. Accordingly, we set $dT = 0$, $dV = 0$ in Helmholtz's ($F$) and Landau's ($\Omega$) thermodynamic potentials:

$$\Omega = F - \hat{\mu}N \quad \text{(S21)}$$

where, in order to link Fröhlich's calculations (Eqs. 11-21 in ref.[29]), $N$ denotes the total excitation number (i.e. phonons) and $\hat{\mu}$ is the associated chemical potential. Now it is matter of some algebraic steps to show that, still for large enough temperature values:

$$\beta\hat{\mu} \approx -\left[(\bar{N} + l^{-1})\frac{\phi_s}{N_z} + \frac{1}{\hbar\beta\varpi}\right]^{-1} \quad \text{(S22)}$$

$\bar{N} \equiv \sum \bar{n}_k$ and $N_z$ being the number of longitudinal electric modes with some characteristic frequency $\varpi$, expected to be nearly independent of $s$. Thus, as:

$$N - \bar{N} \approx \frac{N_z}{\hbar\phi_s\beta\varpi} \quad \text{(S23)}$$

one gets:

$$\hat{\mu} \approx -\frac{N-\bar{N}}{N+l^{-1}}\hbar\varpi \quad \text{(S24)}$$

i.e. $\hat{\mu}$ grows from zero at thermal equilibrium to $\bar{\mu} \approx -\hbar\varpi$ as $N$ increases. As, in this domain, $d(\hat{\mu}N) \approx \bar{\mu}dN$, then $dF \approx \bar{\mu}dN + dW_e$ and:

$$d\Omega = dF - d(\hat{\mu}N) \approx dW_e \quad \text{(S25)}$$

where $W_e$ is the energy influx in response to an applied electric field $\mathbf{E}$, e.g.:

$$\delta W_e = \frac{1}{2}\int(\delta\boldsymbol{E} \cdot \boldsymbol{D})dV \quad \text{(S26)}$$

with displacement vector $\mathbf{D}$ and the integration to be performed inside the dielectric material. Eq. (S25) represents a free energy balance not too far from equilibrium (that may crudely correspond to Eq. S3):

$$\hat{\mu}\frac{dN}{dt} \approx -\frac{dW_e}{dt} + \frac{dF}{dt} \quad \text{(S27)}$$

with $-\delta W_e$ and $-\delta F$ standing respectively for an electric work done on the system and the maximum obtainable work. While, for $\hat{\mu} = 0$, energy influx and outflux equate, when $\hat{\mu} \sim \bar{\mu} < 0$ $(dN > 0)$ the maximum power exerted by the biological matter exceeds that by $W_e$, and condensation may take place.

We remember that, strictly speaking, volume and pressure should not be used as thermodynamic variables in inhomogeneous bodies and/or in the presence of nonuniform or anisotropic (electric) fields. The former remarks thus apply to a thermodynamic system in homogeneous/uniform conditions.

*c.) Large scale–small scale acoustic coupling*

Consider a (reduced) Hamiltonian in second quantization, sum of contributions from excitations of local nature ($a_{\boldsymbol{k}}$) and phonon lattice deformations ($b_{\boldsymbol{q}}$), plus a coupling interaction term ($a_{\boldsymbol{k}}b_{\boldsymbol{q}}$), i.e.:

$$\widehat{H} = H_{\uparrow} + H_{\rightarrow} + H_i \qquad \text{(S28)}$$

given explicitly by:

$$H_{\uparrow} = \sum_{(\boldsymbol{k})} \varepsilon_{\boldsymbol{k}} a_{\boldsymbol{k}}^{\dagger} a_{\boldsymbol{k}}$$

$$H_{\rightarrow} = \sum_{(\boldsymbol{q})} E_{\boldsymbol{q}} b_{\boldsymbol{q}}^{\dagger} b_{\boldsymbol{q}} \qquad \text{(S29)}$$

$$H_i = \sum_{(\mathbf{q},\mathbf{k})} \mu_{\boldsymbol{q}} a_{\mathbf{k}}^{\dagger} a_{\boldsymbol{k}+\boldsymbol{q}} \beta_{\mathbf{q}}$$

where, at wavevectors $\boldsymbol{k}$, $\boldsymbol{q}$, the bosonic operators $(a_{\boldsymbol{k}}^{\dagger}, a_{\boldsymbol{k}})$ and $(b_{\mathbf{q}}^{\dagger}, b_{\boldsymbol{q}})$ create/annihilate the energy quanta $\varepsilon_{\boldsymbol{k}} = \hbar\omega_{\mathbf{k}}$ and $E_{\boldsymbol{q}} = \hbar\Omega_{\mathbf{q}}$ that follow from the respective dispersion laws ($\hbar$ = Planck's constant; $\omega$, $\Omega$ = angular frequencies). The total Hamiltonian $H$ can be derived (except for a constant addendum) by summing Eq. (S28) to the energy-pump Hamiltonian, $H = \widehat{H} + H_P$. However, under non–strong second–order coupling conditions, $H_P$ is shown to commute with both $a_{\mathbf{k}}$ and $b_{\boldsymbol{q}}$ [35] and, therefore, is not regarded in the following. The coupling energy:

$$\mu_{\boldsymbol{q}} = \chi\, \Lambda_{\boldsymbol{q}} \sin^2\left(\frac{\mathbf{q}\cdot\mathbf{a}}{2}\right) \qquad \text{(S30)}$$

includes an anharmonic coupling force of the exciton–phonon type ($\chi > 0$), with values expected to range from fractions to dozens of picometers (pm)[36,37], and the unit cell vector ($\mathbf{a}$), which can be taken to be an average for Bravais lattices that e.g. are not cubic. Unlike the original theories of exciton– or electron–phonon interactions, $\chi$ needs not being normalized to atomic or unit cell numbers[38]. Anyway, when it is non–null, any lattice deformation will impact on the local quasiparticle dynamics (e.g. amplitude or frequency)[39]. Related to a wavevector $\boldsymbol{q}$, the characteristic length:

$$\Lambda_{\boldsymbol{q}} \equiv (\hbar/2M\Omega_{\boldsymbol{q}})^{1/2} \qquad \text{(S31)}$$

accounts instead for the number of atoms involved in the process, with total mass $M$. The last building block of the coupling Hamiltonian is the lattice displacement operator, $\beta_{\boldsymbol{q}} = b_{\boldsymbol{q}} + b_{-\boldsymbol{q}}^{\dagger}$, which normally averages to zero in a phonon number state[40]. The main purpose of this section is to deduce the dynamics of it.

Firstly, we remind the non–trivial commutation rules of bosonic operators adopted here:

$$\left[a_{\boldsymbol{k}}, a_{\boldsymbol{K}}^{\dagger}\right] = \delta_{\boldsymbol{kK}} \quad \left[b_{\boldsymbol{q}}, b_{\boldsymbol{Q}}^{\dagger}\right] = \delta_{\boldsymbol{qQ}} \qquad \text{(S32)}$$

where $(\boldsymbol{k}, \boldsymbol{K})$, $(\boldsymbol{q}, \boldsymbol{Q})$, are pairs of wavevectors, $\delta$ is Kronecker's symbol and, for simplicity, vibrational branch labels are dropped off. Second, the motion equations in time $t$ for the mean displacement operator, $D_{\boldsymbol{q}} \equiv \langle \beta_{\boldsymbol{q}} \rangle = \langle b_{\boldsymbol{q}} \rangle + \langle b_{-\boldsymbol{q}}^{\dagger} \rangle$, follow from combining the quantum Poisson brackets for $b_{\boldsymbol{q}}$, $b_{-\boldsymbol{q}}^{\dagger}$, as they are calculated by the reduced Hamiltonian $\widehat{H}$ applied to any operator $A_{\boldsymbol{u}}$ with wavevector $\boldsymbol{u}$:

$$i\hbar \frac{dA_{\boldsymbol{u}}}{dt} = [A_{\boldsymbol{u}}(t), \widehat{H}]\,, \qquad \text{(S33)}$$

and then from averaging in a (dominant) ground state, $\boldsymbol{q} = \mathbf{g}$, assumed to be coherent with frequency $\Omega_{\mathbf{g}}$. In such a way, one arrives at a steady–state represented by the oscillator law:

$$\left(\frac{\partial^2 D_{\mathbf{g}}}{\partial t^2}\right) + \Omega_{\mathbf{g}}^2 B_{\mathbf{g}} = \frac{2}{\hbar}\,\mu_{-\mathbf{g}}\Omega_{\mathbf{g}} S_{(\mathbf{g})} \qquad \text{(S34)}$$

that is forced by the coupling term on the right side, with $S_{(\boldsymbol{q})} \equiv \sum_{\boldsymbol{k}} \langle a_{\boldsymbol{k}}^{\dagger} a_{\boldsymbol{k}-\mathbf{q}} \rangle$. Getting the solution of Eq. (S34) at any time is generally tough, but we are interested to find the prevailing temporal behaviour. It comes from frequency–transforming ($\mathcal{F}_{\omega}$) Eq. (S34), giving in the end a long–time correlated behaviour of the form:

$$D_{\mathbf{g}}(t) = \frac{\chi}{f_{\mathbf{g}}} \sin^2\left(\frac{\mathbf{g}\cdot\mathbf{a}}{2}\right)\left[\frac{1}{2}F(\Omega_{\mathbf{g}})\cos(\Omega_{\mathbf{g}} t) + 1\right] \qquad \text{(S35)}$$

The first ratio on the right sets the anharmonic coupling in units of the harmonic force $f_{\mathbf{g}} = \hbar\Omega_{\mathbf{g}}/2\Lambda_{\mathbf{g}}$, while the unit term in square brackets corresponds to the asymptotic response of Fourier's transform, $|F(0)| \approx 1$, being generally:

$$F(\omega) = \mathcal{F}_{\omega}\left(\frac{\partial}{\partial t} S_{(\boldsymbol{k})}\right) \qquad \text{(S36)}$$

The net atom displacement $\xi_{\mathbf{g}}^{0}$ now descends from the dimensionless shift in Eq. (S35) that does not average in time to zero, multiplied by the characteristic length scale of the steady state. The latter is simply $\Lambda_{\mathbf{g}}$, calculated by Eq. (S29), the former (say, $D_{\mathbf{g}}^{0}$) is identified by:

$$D_{\mathbf{g}}(t) = D_{\mathbf{g}}^{*}(t) + D_{\mathbf{g}}^{0} \qquad \text{(S37)}$$

with the prerequisite that the temporal mean of $D_{\mathbf{g}}^{*}(t)$ be zero, as fulfilled by Eq. (S35). Therefore, one is left with a classical relation for two parameters, the anharmonic coupling force ($\chi$) and the harmonic spring constant ($k_e$) of lattice atoms:

$$\xi_{\mathbf{g}}^{0} \equiv \left|\Lambda_{\mathbf{g}} D_{\mathbf{g}}^{0}\right| = \frac{\chi}{f_{\mathbf{g}}} \sin^2\left(\frac{\mathbf{g}\cdot\mathbf{a}}{2}\right)\Lambda_{\mathbf{g}} \approx \frac{\chi}{4k_e} \qquad \text{(S38)}$$

as it turns out from replacing the equations for $f_{\mathbf{g}}, \Lambda_{\mathbf{g}}$ and the dispersion law for the acoustic branch:

$$\Omega_{\mathbf{g}}^2 = 4\left(\frac{k_e}{M}\right)\sin^2\left(\frac{\mathbf{g}\cdot\mathbf{a}}{2}\right) \qquad \text{(S39)}$$

Although $k_e$ and $\chi$ could vary along the molecular chain or upon interaction with solvent molecules, Eq. (S38) captures a simple yet fundamental mechanism of protein dynamics, depicted in Figures S5a and S5b and expressible symbolically as:

| *(harmonic spring constant) x (atomic displacement) ~ (coupling work by anharmonic forces)* |
|---|

It is formally well–posed as it writes independently of atom numbers (say $n_N$). To change representation and explicitly introduce $n_N$, we should rescale the typical length, coupling force and Fourier's transform accordingly:

$$\Lambda_{\boldsymbol{q}} \rightarrow \frac{\Lambda_{\boldsymbol{q}}}{\sqrt{n_N}},\ \chi \rightarrow \frac{\chi}{\sqrt{n_N}},\ F(0) \rightarrow\ n_N F(0) \qquad \text{(S40)}$$

but, in the end, Eq. (S38) would not change.

*d.) Notes on atomic displacements upon optical interactions*

In the former analysis local excitations were only let to interact with acoustic phonons. While this remains overall a reliable assumption, contributions of optical nature cannot be ruled out a priori. The study of long-range vibrations in proteins is more complex than in inorganic solids, partly because of dense and nearly featureless dispersion branches, just peaked in the THz range[41]. This would require a dedicated treatment that is far beyond the scope of the present work. What can be achieved here is an estimate of longitudinal optical contributions "per se", based on the view developed so far. To this aim, we introduce an interaction (coupling) Hamiltonian of the form:

$$H_i' = \sum_{(\boldsymbol{q},\boldsymbol{k},\boldsymbol{k'}=\boldsymbol{k}+\boldsymbol{q})} o_{\boldsymbol{q}} a_{\boldsymbol{k}}^{\dagger} a_{\boldsymbol{k'}} \beta_{\boldsymbol{q}} \qquad \text{(S41)}$$

with $|o_{\boldsymbol{q}}| = (q_e/|\boldsymbol{q}|a)\sqrt{2\pi\hbar\nu_{\boldsymbol{q}}/na\epsilon^*}$ [38], $q_e$ denoting the electron charge, $\nu_{\boldsymbol{q}}$ the optical frequency with wavenumber $\boldsymbol{q}$, and $1/\epsilon^* = 1/\epsilon_\infty - 1/\epsilon_S$ being the difference of the reciprocals of the high-frequency (visible/infrared) and static dielectric permittivities. An effective cell volume $na^3$ then can be defined as a function of the implied chain step (or lattice constant) $a$ and an average natural number $n$, measure of periodicity. Nonphysical effects from unshielded Coulomb potentials at very small $|\boldsymbol{q}|$ here should be disregarded, as real molecules exhibit electric screening. An estimation of the optical contribution $\xi_{\mathbf{g}}^{O}$ to the mean atomic shift in a ground steady state ($\boldsymbol{q} = \mathbf{g}$) stems by approximating $\nu_{\boldsymbol{q}} \sim \Omega_{\boldsymbol{q}}$. Replacing the acoustic coupling in Eq. (S28) with Eq. (S41), one finds that:

$$\xi_{\mathbf{g}}^{O} \approx \frac{q_e \lambda_F^2}{2\pi\sqrt{\pi n \nu k_e \epsilon^*}} \qquad \text{(S42)}$$

is still a classical relationship, where $\lambda_F \approx 2\pi/|\boldsymbol{q}|$ is the spatial wavelength.

The quantity $\epsilon^*$ can be evaluated by a mean field model for a dielectric mixture (protein *p*, water *w*), for it is known that the overall dielectric constant in the longitudinal direction roughly descends from $\epsilon = \varphi_p \epsilon^p + (1 - \varphi_p)\epsilon^w$ ($\varphi_p$ = protein volume fraction)[42]. Dielectric features of proteins are tough to be approached quantitatively, they can sensibly change upon structural rearrangements and ionization phenomena [43]. However reasonable values of the pure protein phase should stay in $\epsilon_S^p \sim (2 - 4)$ [43], which are expected to decrease to $\epsilon_\infty^P \sim (1.5 - 3)$ at frequencies $\gtrsim 0.5$ THz[44]. Degrees of freedom of electric dipoles of water molecules are affected by the crystal lattice, interfacial order perturbations and nanoconfinement[45]. As specific information on the hydration structure of trypsin molecules is lacking, a range $\epsilon_S^w \sim (30 - 50)$ may be assumed as a representative dielectric reduction throughout the sample with $\epsilon_\infty^w \sim (4 - 6.5)$ at room temperature[45,46]. Weighting such values with $\varphi_w \equiv 1 - \varphi_p \sim (0.4 - 0.5)$ [34], it turns out $1/\epsilon^* = 1/\overline{\epsilon}_\infty - 1/\overline{\epsilon}_S \sim (0.1 - 0.4)$.

When the chain step stands for the size of a unit cell of volume $v \approx a^3$, we get $a \sim 3.75$ nm per trypsin molecule and $\xi_{\mathbf{g}}^O \le 0.1$ pm ($n \ge 1$) with $\lambda_F \sim (1 - 10)$ µm. If it goes down to more typical scales of H–bonds or dipolar chain steps, i.e. $a \sim (2 - 5)$ Å [39,47,48], then $\xi_{\boldsymbol{g}}^P \le 0.3$ pm ($n \ge 10^3$) in the identical $\lambda_F$ range. Occurrence of a micrometer correlation length is definutely reasonable in a biological process. To conclude, as compared to the acoustic case, we expect possible optical displacements to be smaller by at least one order of magnitude.

*e.) Including the local excitation dynamics*

The former analysis was focused on a coherent steady state in the lattice deformation dynamics. A similar calculation can be developed for the joint oscillatory motion of large–scale and smaller– scale excitations. To Heisenberg's equations for lattice modes, one now is required to add the dynamics for local quasiparticles by means of Eq. (S33), for example, $i\hbar\dot{a}_{\boldsymbol{k}}^\dagger(t) = [a_{\boldsymbol{k}}^\dagger(t), \widehat{H}]$. This will give rise to a dual "entrained" motion of phonon–like oscillators and vibrations that are more local in nature. It suffices to introduce the displacement operator, $\alpha_{\boldsymbol{k}} = a_{\boldsymbol{k}} + a_{-\boldsymbol{k}}^\dagger$, with characteristic length $\lambda_{\mathbf{k}} \equiv \sqrt{\hbar/(2m\omega_{\mathbf{k}})}$ and mass ($m$) scale of the involved amino acidic residues. Unlike the self-focusing in the frequency spectra of coherent phonon states, with rather localised interactions in $\boldsymbol{q}$–space, we attempt an energy dispersion law in $\boldsymbol{k}$–space to be in the first order as reasonably constant, $\omega_{\boldsymbol{k}} \approx \omega_a = \varepsilon_a/\hbar$ [38,49 38,49].

To discern each term carefully, one can insert in the Hamiltonian the interaction terms between wavevectors $\boldsymbol{k}$ and $\boldsymbol{k} + \boldsymbol{h}$ (i.e. $a_{\boldsymbol{k}}^\dagger a_{\boldsymbol{k}+\boldsymbol{h}}$, $\boldsymbol{h}$ fixed), then express the motion equations for $a_{\boldsymbol{k}}, a_{-\boldsymbol{k}}^\dagger$ and $\alpha_{\boldsymbol{k}} \equiv a_{\boldsymbol{k}} + a_{-\boldsymbol{k}}^\dagger$. The result is:

$$\hbar^2\left(\frac{\partial^2\alpha_k}{\partial t^2}\right) = -\,\varepsilon_k\big(\varepsilon_k + \sum_q \mu_q\beta_q\big)\alpha_k - \varepsilon_{k+h}\alpha_{k+h}\sum_q \mu_q\beta_q + \hat{\alpha}^{\dagger}_{k+h}\sum_q \hbar\Omega_q\mu_q\hat{\beta}^{\dagger}_q - \alpha_{k+h}\sum_{qq'}\mu_q\mu_{q'}\beta_q\beta_{q'} \quad \text{(S43)}$$

all symbols being still as in main text, obviously with $\mu_q = \mu_{-q}$, $\omega_{\mathbf{k}} = \omega_{-\mathbf{k}}$, $\Omega_{\mathbf{q}} = \Omega_{-\mathbf{q}}$. Hatted operators connect to their conjugates (momentum), $p_k = -i\sqrt{m\hbar\omega_k/2}\,\hat{\alpha}_{-k}$, $P_q = -i\sqrt{m\hbar\Omega_q/2}\,\hat{\beta}_{-q}$. Next steps come from resuming $\boldsymbol{h} \to \mathbf{0}$, dealing with a ground–state, $\boldsymbol{q} \to \mathbf{g}$, and neglecting terms at second-order in $\beta_q$. Taking the average of Eq. (S43), $A_k \equiv \langle\alpha_k\rangle$, returns in this case that:

$$\hbar^2\left(\frac{\partial^2 A_k}{\partial t^2}\right) = -\varepsilon_k^2 A_k - \mu_g\big(2\varepsilon_k\langle\alpha_k\beta_{\mathbf{g}}\rangle + \hbar\Omega_g\langle\hat{\alpha}_k\hat{\beta}_{\mathbf{g}}\rangle\big) + \mathcal{O}(\beta^2) \qquad \text{(S44)}$$

This expansion is acceptable, second-order terms in $\beta_{\mathbf{g}}$ predict lattice perturbations $\sim 10^{-2}$ smaller than $\alpha_k\beta_{\mathbf{g}}$ contributions. On assuming a delocalized coupling, $\langle\alpha_{\mathbf{k}}\beta_{\mathbf{g}}\rangle \sim AB_{\mathbf{g}}$, and noting, from the mechanical momentum definition, that $\langle\hat{\alpha}_{\mathbf{k}}\hat{\beta}_{\mathbf{g}}\rangle \sim -\,\dot{A}\dot{B}_{\mathbf{g}}/(\omega_a\Omega_g)$, one finally obtains a second dynamic law:

$$\frac{\partial^2 A}{\partial t^2} + \omega_a^2 A = -2\mathrm{w}_{\mathbf{g}}\left[2\omega_a A D_{\mathbf{g}} - \frac{1}{\omega_a}\left(\frac{\partial A}{\partial t}\right)\left(\frac{\partial D_{\mathbf{g}}}{\partial t}\right)\right] \qquad \text{(S45)}$$

where, besides $\Omega_{\mathbf{g}}$, two other frequencies take place, $\omega_a$ and $\mathrm{w}_{\mathbf{g}} = \mu_{\mathbf{g}}/\hbar$. This brings the joint equation system, Eq. (S34) and Eq. (S45), to take the following (dimensionless) form:

$$\left(\frac{\partial^2 D_{\mathbf{g}}}{\partial\tau^2}\right) = -\,D_{\mathbf{g}} - \gamma_{\mathrm{g}}(A^2+1)$$

$$\left(\frac{\partial^2 A}{\partial\tau^2}\right) = -\,\frac{1}{4}\gamma_a^2 A - \gamma_{\mathrm{g}}\left[\gamma_a A D_{\mathbf{g}} - \frac{\gamma_{\mathrm{g}}}{\gamma_a}\left(\frac{\partial A}{\partial\tau}\right)\left(\frac{\partial D_{\mathbf{g}}}{\partial\tau}\right)\right] \qquad \text{(S46)}$$

with $\tau = \Omega_{\mathbf{g}}t$, $\gamma_{\mathrm{g}} = 2\mathrm{w}_{\mathbf{g}}/\Omega_{\mathbf{g}}$, $\gamma_a = 2\omega_a/\Omega_{\mathbf{g}}$. In the forthcoming subsection, we concentrate on the calculation of displacements in the steady–states belonging to Eqs. (S46).

*f.) Steady–state analysis of atomic shifts in the entrained system*

Eqs. (S46) allow to evaluate the atomic displacements ($\xi$) in two protein subsystems: a lattice of atoms hosting large–scale phonon deformations ($\xi_{\mathbf{g}}$); a second domain with local vibrational excitations coupled to the lattice ($\xi_a$). In this 2–dimensional displacement space, Eqs. (S46) has three fixed points (or steady–states) of the general form, $\vec{P_i} \in (A \times D_{\mathbf{g}})_i$ ($i = 0, 1, 2$), i.e.:

$$\vec{P}_0 = [0, -\gamma_{\mathrm{g}}],\ \vec{P}_{1,2} = \left[\pm\big(\gamma_a/4\gamma_{\mathrm{g}}^2 - 1\big)^{\frac{1}{2}}, -\gamma_a/4\gamma_{\mathrm{g}}\right] \qquad \text{(S47)}$$

associated with a secular polynomial of the fourth degree. It can be demonstrated that purely imaginary eigenvalues (i.e. oscillatory solutions) for $\vec{P}_0$ need $\gamma_{\mathrm{a}} > 4\gamma_{\mathrm{g}}^2$, outside which two of them are zero or real positive (unstable). On the contrary, for $\vec{P}_{1,2}$ to be oscillatory, the condition $1 > 8\gamma_a\gamma_{\mathrm{g}}^2 - 2\gamma_a^2 \geq 0$ is to be met. However, Eqs. (S46) are mostly meant here to show that including local quasiparticles in the study of lattice deformations can afford better predictions for atomic shifts. Their thorough dynamic analysis falls outside the aims of the present work.

To get new estimates in this setting, we multiply again the absolute values of their coordinates by the respective typical lengths, $\lambda_a$ and $\Lambda_{\mathbf{g}}$, yielding displacement vectors of the form $|\vec{\xi}_i| = (\xi_{ai}, \xi_{\mathbf{g}i})$, being $\xi_{ai} = |A_i\lambda_a|$, $\xi_{\mathbf{g}i} = |D_{\mathbf{g}i}\Lambda_{\mathbf{g}}|$. Replacing the definitions of $\gamma_a$, $\gamma_{\mathbf{g}}$ and carrying out some algebra lead to the following formulas, set in the explicit physico–chemical properties of the protein molecule:

$$|\vec{\xi}_0| = \left(0, \frac{\chi}{4k_e}\right)$$

$$\frac{1}{\hbar}|\vec{\xi}_{1,2}| = \left(\left|\frac{1}{m}\left(\frac{2k_e^2}{M\chi^2\Omega_{\mathbf{g}}^2}\right) - \frac{1}{2\hbar\omega_a}\right|^{\frac{1}{2}}, \frac{k_e\omega_a}{\chi M\Omega_{\mathbf{g}}^2}\right) \qquad \text{(S48)}$$

The lattice component of the (classical) ground displacement vector (0) is $\xi_{\mathbf{g}} = \xi_{\mathbf{g}}^0$ (see main text, Eq. 3), with zero dipole–like component. This recovers the phononic steady–state without local excitations, and is a consequence of the factor $-\gamma_{\mathrm{g}} A^2$ in the first of Eqs. (S46). Vectors 1,2 extend the former steady–state into a dual dynamic picture with $\xi_a \neq 0$ and own the same absolute shift. Eqs. (S48) suit to get the numerical evaluations in the main text but are fully equivalent to their energy representation in Eqs. (4).

*g.) Collective Fluctuations and B-factors ($B$)*

The transition from a phonon number state to a coherent state, in principle, should go hand in hand with a reduction of fluctuations (an increase in the molecular order)[40]. However, the trend of B-factors along the protein chain behaves quite irregularly in our experiment, even though atomic shifts exhibit significant correlations. It seems that a certain energy redistribution takes place, with fractions of atoms fluctuating less or more around the control values. We now verify that the average B-factor $\langle B \rangle$ may even increase upon the establishment of a meaningful coupling between lattice deformations and more local excitations. This is not incompatible with a large–scale dynamic process exhibiting coherence. Despite the protein molecule as a whole cannot be regarded as it were in a fully coherent state, yet concerted reductions of *B*-factors can develop along particular chain pathways.

Here we concentrate on B-factors of local quasiparticles interacting with phonons of frequency $\Omega_{\mathbf{g}}$. The simplest way to proceed, according to the theory of canonical transformations, is working with the energy eigenvalues of the (reduced) Hamiltonian. For an ensemble of local excitations, the Hamilton function in diagonal form, $\widehat{H} \to \widehat{\mathcal{H}}$, simplifies into[38]:

$$\widehat{\mathcal{H}} \; = \; E_A \sum_{\boldsymbol{k}} \bar{a}_{\boldsymbol{k}}^{\dagger} \bar{a}_{\boldsymbol{k}} \; + \; \hbar\Omega_{\boldsymbol{g}} \sum_{\boldsymbol{k}} \bar{b}_{\boldsymbol{k}}^{\dagger} \bar{b}_{\boldsymbol{k}} \tag{S49}$$

that, strictly speaking, holds for coupling perturbations smaller than two-phonon energy thresholds, i.e. $E_A < \hbar\omega_a + 2\Omega_{\boldsymbol{g}}$. Symbols, again, are as in the main text, overlines denoting the operators canonically transformed by the anti–Hermitian coupling Hamiltonian, i.e. $H_i \to \mathcal{H}_i$ [50], whereas:

$$E_A \; = \; \hbar\omega_a \; - \; \frac{1}{n\hbar\Omega_{\boldsymbol{g}}} \sum_{\boldsymbol{q}} \mu_{\boldsymbol{q}}^2 \tag{S50}$$

We are interested to see how the perturbation to the former energy eigenvalue affects, on the average, the measurements of $B$. An effective way to deal with a stochastic process parametrized by a number of $n$ atomic/molecular constituents (the dimensionality of a sample of amino acids, dipoles, etc. producing the local excitations) is to rearrange Eq. (S50) by letting the energy $\mu_{\boldsymbol{q}}$ to behave as a random variable:

$$E_A = \hbar\omega_a - \langle \mu^2 \rangle - \frac{\sigma_\mu}{\sqrt{n}\hbar\Omega_{\boldsymbol{g}}} \, E_n \tag{S51}$$

$E_n$ converging in law e.g. to a Gaussian distribution with zero mean, $E_n \longrightarrow N(0, \sigma_\mu)$ :

$$E_n = \frac{1}{\sqrt{n}\sigma_\mu} \left[ \sum_{i=1}^{n} \mu_{\boldsymbol{q}_i}^2 - n\langle \mu^2 \rangle \right] \tag{S52}$$

The calculation of $\langle \mu^2 \rangle = \left\langle \psi_c \middle| \mu_{\boldsymbol{q}}^2 \middle| \psi_c \right\rangle$ can be carried out by a coherent–like probability amplitude[51]:

$$\psi_c^* \psi_c(q) = \left( \pi\hbar\bar{M}\Omega_{\boldsymbol{g}} \right)^{-\frac{3}{2}} e^{-\hbar|\boldsymbol{q}|^2/\bar{M}\Omega_{\boldsymbol{g}}} \tag{S53}$$

$\bar{M}$ now denoting the atomic mass the B-factor refers to. To fix the ideas we imagine that $\langle B \rangle$ moves from the value in the unperturbed state (*off*) to a new value (*on* state) set by the coupling interaction. A shift is implied by Eq. (S51) as $\Delta B = B_{on} - B_{off} > 0$. Let $\theta \equiv n\langle \mu^2 \rangle / \left( k_B \hbar\Omega_{\boldsymbol{g}} \right)$ be the temperature variation produced by the coupling term $\langle \mu^2 \rangle$, it turns out in the long wavelength limit that:

$$\theta \; \approx \; \frac{\pi^3}{3} \frac{n\chi^2}{k_{\mathrm{e}}} \left( \frac{a}{\lambda_F} \right)^{\frac{1}{2}} \left( \frac{\Lambda_h}{\lambda_F} \right)^3 \tag{S54}$$

with typical length scale $\Lambda_h = (4\pi^2\hbar^2/k_{\mathrm{e}}\bar{M})^{\frac{1}{4}}$. We can now afford a theoretical first–order estimate (*th*) for the B-factor change, $(\Delta B/B_0)_{th} \approx 2\theta/\Theta$, with $B_0 \equiv B_{off}$ and $\Theta$ is an effective Debye's temperature.

This view holds well for a crystal of harmonic oscillators, their density of vibrational states being proportional to the square of frequency when the normal mode wavelengths are much longer than the atomic spacing. Despite a protein crystal displays some amorphousness degree, coupling anharmonically the low-frequency modes[52], Eq. (S54) yields a reasonable estimate to be compared with some typical experimental magnitudes. Adopting $B_0 \sim (17-18)$ Å², $\delta B \sim +(4-5)\cdot 10^2$ Å², $\Theta = \hbar\omega_a/k_B \sim (5-100)$ K, $n \sim (10^2 - 10^7)$ and the volume of the trypsin unit cell, $v \sim a^3$, a relative experimental (*ex*) shift amounting to $(\delta B/B_0)_{ex} \sim (2-3)\cdot 10^{-3}$ is theoretically recovered in an atomic population of C, O, N atoms ($\bar{M} = 12-16$ amu) when $\lambda_F \sim (2-20)$ nm. Wavelengths tend clearly to increase with increasing dipole number, e.g. $\lambda_F \sim (1-20)$ µm if $n \sim 10^{12}-10^{14}$. It is also important to remember that a statistically meaningful shift in the B-factor distribution may not always take place, and should be verified with the tools of inferential statistics on a case–by–case basis.

*h.) Squeezed-like fluctuations in B-factors*

The first time derivative of $B_{\boldsymbol{q}} \equiv \beta_{\boldsymbol{q}}\beta_{-\boldsymbol{q}}^{\dagger}$ identifies a two-mode quadrature squeezing operator[53]:

$$-\frac{i}{2\Omega_{\boldsymbol{q}}}\left(\frac{\partial B_{\boldsymbol{q}}}{\partial t}\right) = b_{\boldsymbol{q}}^{\dagger}b_{-\boldsymbol{q}}^{\dagger} - b_{-\boldsymbol{q}}b_{\boldsymbol{q}} \tag{S55}$$

returning a class of forced oscillator laws given by:

$$\frac{i}{4\Omega_{\boldsymbol{q}}^2}\left(\frac{\partial^2 B_{\boldsymbol{q}}}{\partial t^2}\right) + B_{\boldsymbol{q}} = b_{-\boldsymbol{q}}^{\dagger}b_{-\boldsymbol{q}} + b_{\boldsymbol{q}}b_{\boldsymbol{q}}^{\dagger} - \frac{1}{2}\frac{w_{\boldsymbol{q}}}{\Omega_{\boldsymbol{q}}}\left(\hat{P}_{\boldsymbol{q}} + \hat{P}_{-\boldsymbol{q}}\right)S_{(\boldsymbol{k})} \tag{S56}$$

Here, $\hat{P}_{\boldsymbol{q}} = i(b_{\boldsymbol{q}}^{\dagger} - b_{-\boldsymbol{q}})$ is the operator for the generalized $\boldsymbol{q}$–impulse, and every symbol is defined as in the main text. The dependence of $B_{\boldsymbol{q}}$ on $\hat{P}_{\boldsymbol{q}}$ is a further indication of the influence of the kinematic state of the protein on atomic fluctuations. The previous equation is symbolically resolved by exploiting the solutions for bosonic operators and their Hermitian conjugates, e.g.:

$$b_{\boldsymbol{q}} = -\frac{i}{\hbar}\mu_{\boldsymbol{q}}e^{-i\Omega_{\boldsymbol{q}}t}\int_{-\infty}^{t} S_{(\boldsymbol{k})}(t')e^{i\Omega_{\boldsymbol{q}}t'}dt + c_{\boldsymbol{q}} \tag{S57}$$

$c_{\boldsymbol{q}}$ being is the c–number carrying the chosen boundary condition. Accordingly:

$$\frac{\partial^2 B_{\boldsymbol{q}}}{\partial\tau_q^2} + B_{\boldsymbol{q}} = B_n + \frac{1}{2}\gamma_q^2\int_{-\infty}^{\tau_q}\langle S_{(\boldsymbol{k})}\rangle e^{-\frac{i}{2}\tau'_q}d\tau'_q\int_{-\infty}^{\tau_q}\langle S_{(\boldsymbol{k})}\rangle e^{\frac{i}{2}\tau'_q}d\tau'_q$$

$$-\frac{i}{2}\gamma_q^2\left[e^{\frac{i}{2}\tau_q}\int_{-\infty}^{\tau_q}\langle S_{(\boldsymbol{k})}\rangle e^{-\frac{i}{2}\tau'_q}d\tau'_q + e^{-\frac{i}{2}\tau_q}\int_{-\infty}^{\tau_q}\langle S_{(\boldsymbol{k})}\rangle e^{\frac{i}{2}\tau'_q}d\tau'_q\right] \tag{S58}$$

where $\tau_q \equiv 2\Omega_{\boldsymbol{q}}$t, $\gamma_q \equiv \mathrm{w}_{\boldsymbol{q}}/\Omega_{\boldsymbol{q}} = (\chi/4k_e)\sqrt{M\Omega_{\boldsymbol{q}}/2\hbar}$ and $B_n$ gets the c-number to a proper integration constant. To derive the dominant time–response for $B_{\boldsymbol{q}} = B_{\boldsymbol{q}}(t)$, the next SI (3i) Fourier–transforms Eq. (S58). The final result, at third order in $\gamma_q$, is:

$$B_{\boldsymbol{q}}(\tau_q) \approx B_n + \frac{i}{3}\langle s\rangle\gamma_q^3\left[8\cos(\tfrac{1}{2}\tau_q) - \cos(\tau_q) - \cos(\tfrac{1}{2}\gamma_q\tau_q)\right] + O(\gamma_q^4) \quad \text{(S59)}$$

$\langle s\rangle$ being a mean proportionality coefficient in Fourier's transform of $S_{(\boldsymbol{q})}$, here set to be representative of a Brownian oscillator system [54]. The previous relation asserts that a nonlinear B-factor modulation develops in time (accompanied by a coherent atomic shift, see Figure S6).

*i.) Details of direct and inverse Fourier transforms*

In real physical units, the Fourier transform of Eq. (S58) is:

$$(4\Omega_{\boldsymbol{q}}^2 - \omega^2)\mathcal{B}_{\boldsymbol{q}}(\omega) = 8\pi^2\Omega_{\boldsymbol{q}}^2 B_n\delta(\omega) - 4i\Omega_{\boldsymbol{q}}\mathrm{w}_{\mathbf{q}}^2\Big\{-i\left(\tfrac{1}{\omega-\Omega_{\boldsymbol{q}}} + \tfrac{1}{\omega+\Omega_{\boldsymbol{q}}}\right)\mathcal{S}_\omega(\omega) + \pi\big[\mathcal{S}_\omega(\Omega_{\boldsymbol{q}})\delta(\omega+\Omega_{\boldsymbol{q}}) + \mathcal{S}_\omega(-\Omega_{\boldsymbol{q}})\delta(\omega-\Omega_{\boldsymbol{q}})\big]\Big\} + 4\Omega_{\boldsymbol{q}}^2\mathrm{w}_{\boldsymbol{q}}^2\Big\{2\pi^2\mathcal{S}_\omega(\Omega_{\boldsymbol{q}})\mathcal{S}_\omega(-\Omega_{\boldsymbol{q}})\delta(\omega) - \tfrac{i}{\omega}\big[\mathcal{S}_\omega(-\Omega_{\boldsymbol{q}})\mathcal{S}_\omega(\omega-\Omega_{\boldsymbol{q}}) + \mathcal{S}_\omega(\Omega_{\boldsymbol{q}})\mathcal{S}_\omega(\omega+\Omega_{\boldsymbol{q}})\big] - \tfrac{1}{\pi}\mathcal{I}(\Omega_{\boldsymbol{q}})\Big\} \quad \text{(S60)}$$

$\mathcal{B}_{\boldsymbol{q}}$ and $\mathcal{S}_\omega$ being the Fourier transforms of $B_{\boldsymbol{q}}$ and $S_{(\boldsymbol{q})}$, while:

$$\mathcal{I}(\Omega_{\boldsymbol{q}}) = P.V.\int_{-\infty}^{\infty}\frac{\mathcal{S}_\omega(y-\Omega_{\boldsymbol{q}})\mathcal{S}_\omega(\omega-y-\Omega_{\boldsymbol{q}})}{y(\omega-y)}dy \quad \text{(S61)}$$

denotes the principle value integral of a convolution function. To explicit the Fourier response for $\mathcal{S}_\omega$, one can resort to the Brownian oscillator model[54], centred for simplicity in the characteristic coupling frequency ($\pm\mathrm{w}_{\boldsymbol{q}}$):

$$\mathcal{S}(\Omega_{\boldsymbol{q}}) = \frac{\langle s\rangle}{W_{\boldsymbol{q}}^2 + \left(\frac{\phi}{2} - i\omega\right)^2} \quad \text{(S62)}$$

where $\phi$ and $\langle s\rangle$ are respectively a damping constant and a phenomenological coefficient related to the inverse of the reduced mass, with $W_{\boldsymbol{q}}^2 \equiv \mathrm{w}_{\boldsymbol{q}}^2 - \phi^2/4$. Eq. (S62) interprets the atomic nuclear motions as harmonic oscillators fulfilling Langevin's equation, and was successfully applied to low-frequency protein spectra ($\leq$ 6 THz)[55]. In line with our analyses, we focus on the limit of negligible damping ($\phi \ll 1$), for $\mathcal{S}_\omega(\omega) = \mathcal{S}_\omega(-\omega)$. It is therefore possible to single out four general contributions to $\mathcal{B}_{\boldsymbol{q}}(\omega)$ to be anti–transformed in the temporal domain. Alongside, the principal integral in Eq. (S61) vanishes. Let:

$$1 - \frac{\omega^2}{(2\Omega_{\boldsymbol{q}})^2} \equiv \frac{1}{O_{\boldsymbol{q}}(\omega)}$$

$$\left(1 - \frac{\Omega_{\boldsymbol{q}}^2}{\mathrm{w}_{\boldsymbol{q}}^2}\right)^2 \equiv \frac{1}{P_{\boldsymbol{q}}}$$

they are:

$$2\pi O_{\boldsymbol{q}}\big(B_n + \pi s^2 P_{\boldsymbol{q}}\big)\delta(\omega) \;\rightarrow\; B_n + \pi s^2 P_{\boldsymbol{q}}$$

$$-i\pi s \frac{\mathrm{w}_q}{\Omega_q}\sqrt{P_{\boldsymbol{q}}}\,O_{\boldsymbol{q}}\big[\delta\big(\omega + \Omega_{\boldsymbol{q}}\big) + \delta\big(\omega - \Omega_{\boldsymbol{q}}\big)\big] \;\rightarrow\; -\frac{4is}{3}\frac{\mathrm{w}_q}{\Omega_q}\sqrt{P_{\boldsymbol{q}}}\cos(\Omega_{\boldsymbol{q}}t) \qquad \text{(S63)}$$

$$2sO_{\boldsymbol{q}}\frac{\mathrm{w}_{\boldsymbol{q}}^3\omega}{\Omega_{\boldsymbol{q}}\big(\omega^2-\Omega_{\boldsymbol{q}}^2\big)\big(\omega^2-\mathrm{w}_{\boldsymbol{q}}^2\big)} \;\rightarrow\; -\frac{i}{6\Omega_{\boldsymbol{q}}^2}\left[-\frac{\cos\big(\Omega_{\boldsymbol{q}}t\big)}{\Omega_{\boldsymbol{q}}^2-\mathrm{w}_{\boldsymbol{q}}^2}+\frac{\cos\big(2\Omega_{\boldsymbol{q}}t\big)}{4\Omega_{\boldsymbol{q}}^2-\mathrm{w}_{\boldsymbol{q}}^2}+\frac{3\cos\big(\mathrm{w}_{\boldsymbol{q}}t\big)}{4\Omega_{\boldsymbol{q}}^2-5\mathrm{w}_{\boldsymbol{q}}^2+\left(\frac{w_{\boldsymbol{q}}^2}{\Omega_{\boldsymbol{q}}}\right)^2}\right]$$

$$-\frac{2is^2}{\omega}\sqrt{P_{\boldsymbol{q}}O_{\boldsymbol{q}}}\frac{\mathrm{w}_{\boldsymbol{q}}^2\big(\mathrm{w}_{\boldsymbol{q}}^2-2\omega\Omega_{\boldsymbol{q}}\big)}{\mathrm{w}_{\boldsymbol{q}}^4-4\mathrm{w}_{\boldsymbol{q}}^2\Omega_{\boldsymbol{q}}\omega-\big(\omega^2-\Omega_{\boldsymbol{q}}^2\big)^2} \;\rightarrow\; \approx\; 4is^2\left(\frac{\mathrm{w}_{\boldsymbol{q}}}{\Omega_{\boldsymbol{q}}}\right)^4\sin\big(2\Omega_{\boldsymbol{q}}t\big)$$

By Taylor–expanding each contribution in $\gamma_q$, one arrives at the time–dependent expression given by:

$$\delta B_{\boldsymbol{q}}(t) \;\approx\; \frac{i}{3}s\gamma_q^3\big[8\cos\big(\Omega_{\boldsymbol{q}}t\big)-\cos\big(2\Omega_{\boldsymbol{q}}t\big)-3\cos\big(\mathrm{w}_{\boldsymbol{q}}t\big)\big]+4is^2\gamma_q^4\sin\big(2\Omega_{\boldsymbol{q}}t\big) \qquad \text{(S64)}$$

where $\delta B_{\boldsymbol{q}}(t) \equiv B_{\boldsymbol{q}}(t) - B_n$. To proceed we stick to Eq. (S59) at third order in $\gamma_q = (\chi/4k_e)\sqrt{M\Omega_{\boldsymbol{q}}/2\hbar}$, with $\tau_q \equiv 2\Omega_{\boldsymbol{q}}t$, which holds well especially for $(n_N\,\Omega_{\boldsymbol{q}}) \lesssim 10^2$ (amu THz). Including higher orders in $\gamma_q$ will obviously enlarge its applicability domain.

*l.) Evaluation of the final B-factor*

To pass from Eq. (S59) to a measurable B-factor, one should build up the usual sum over normal modes ($\boldsymbol{q}$) and vibrational branches ($j$). The most precise method would be starting from Debye-Waller's factor, inclusive of explicit scalar products of the diffraction vector with normalized eigenvectors, giving the displacement direction of each atom from its crystallographic position[38]. Nevertheless, as we are mainly focused on predicting the magnitude of B-factors and have not enough analytic knowledge of the involved polarization vectors, we will limit ourselves to summing up Eq. (S59) in a Debye-like continuum, for:

$$\delta B \equiv \sum_{\boldsymbol{q},j}\delta B_{\boldsymbol{q}j}(t) = \frac{i}{8\pi^2}\sqrt{\frac{m}{2\hbar}}\langle s\rangle\left(\frac{a\xi_g^0}{\bar{c}}\right)^3 I_{5/2}(\Omega_M,t) \qquad \text{(S65)}$$

depends on some cut–off frequency $\Omega_M$ and the sound speed $\bar{c}$, averaged over longitudinal ($j = L$) and two transverse ($j = T_1, T_2$) acoustic branches. The final temporal behaviour descends from:

$$I_{5/2}(\Omega_M,t) = \int_0^{\Omega_M}\Omega^{\frac{5}{2}}G(\Omega,t)d\Omega \qquad \text{(S66)}$$

with:

$$G(\Omega, t) = 8\cos(\Omega t) - \cos(2\Omega t) - 3\cos\left(\Omega^{3/2}\zeta t\right) \tag{S67}$$

and $\zeta = \xi_{\boldsymbol{g}}^0\sqrt{M/2\hbar}$. Evaluating this integral analytically requires the use of special tools – such as optical Fresnel integrals, incomplete gamma and error functions – but a suitable approximation can be inferred from the dominant terms arising, at long times, from each of the three cosine contributions to $G$, i.e.:

$$I_{5/2}^{+}(\Omega_M, t) \sim \frac{\Omega_M^{\frac{5}{2}}}{t}\left[8\sin(\Omega_M t) - \frac{1}{2}\sin(2\Omega_M t)\right] - \frac{2\Omega_M^2}{\zeta t}\sin\left(\Omega_M^{\frac{3}{2}}\zeta t\right) \tag{S68}$$

Accordingly, as $\bar{c}^3 = (a\Omega_M)^3/(6\pi^2)$ [38], one is left with:

$$\delta B\ (t) \sim \frac{3i}{4}\sqrt{\frac{m}{2\hbar}}\langle s\rangle\left(\frac{\xi_g^0}{\Omega_M}\right)^3 I_{5/2}^{+}(\Omega_M, t) \tag{S69}$$

which is finally rearranged to give:

$$\delta B\ (t) \sim \frac{3n_N i\langle s\rangle}{2\Omega_M t}\left(\xi_{\boldsymbol{g}}^0\right)^2\left\{\zeta\sqrt{\Omega_M}\left[4\sin(\Omega_M t) - \frac{1}{4}\sin(2\Omega_M t)\right] - \sin\left(\Omega_M^{\frac{3}{2}}\zeta t\right)\right\} \tag{S70}$$

The two decaying factors in front of the periodic functions on the right identify the B-factor amplitudes discussed in the main text upon $\langle s\rangle \sim n_N$ (Eqs. 4). The first, classical in nature, is the front factor of Eq. (S70); the second is semiclassical/quantum and given by the same term multiplied by $\zeta\sqrt{\Omega_M}$. A concrete realization of Eq. (S70) is illustrated in Figure S7.

**Table S1: Comparison of crystallographic quality indicators.** The mean and standard deviation are displayed. The values in parenthesis refer to the reflections in the highest resolution bin. The integration involved multiple rotation wedges which were indexed separately.

| **Experimental parameters** | **$DI_{off}$** | **$DI_{THz}$** |
|---|---|---|
| Wavelength (Å) | 1.112 | |
| Bandwidth | 0.01 | |
| Detector distance (mm) | 142 | |
| Rotation step/oscillation range (°) | 0.1 | |
| Mean flux (photons $s^{-1}$) | $2 \times 10^7$ | |
| Exposure time per frame | 150 fs | |
| Photons per dataset | $1.3 \times 10^{12}$ | |
| Approximate collection time | 18 h | |
| **Data reduction statistics** | **$F_{off,n}$** | **$F_{THz,n}$** |
| Resolution range (Å) | 6.44 - 1.44 (1.48 - 1.44) | 6.44 - 1.44 (1.48 - 1.44) |
| Space group | $P2_12_12_1$ | $P2_12_12_1$ |
| Unit cell A length (Å) | 55.140±0.146 | 55.139±0.144 |
| Unit cell B length (Å) | 58.888±0.153 | 58.888±0.154 |
| Unit cell C length (Å) | 67.915±0.409 | 67.929±0.417 |
| Total number of reflections | 87225 ±311 (361±2) | 86982±343 (362±2) |
| Number of unique reflections | 32810±28 (329±2) | 32780±35 (328±2) |
| Multiplicity | 2.7±0.0 (1.6±0.0) | 2.7±0.0 (1.6±0.0) |
| Completeness (%) | 80.0±0.1 (11.1±0.1) | 80.0±0.1 (11.1±0.1) |
| $CC_{1/2}$ (%) | 97.9±0.9 (40.3±12.0) | 97.9±0.9 (28.8±15.0) |
| $\langle I/\sigma(I)\rangle$ | 6.00±0.16 (0.52±0.04) | 5.90±0.18 (0.55±0.05) |
| $R_{merge}$ (%) | 12.5±0.6 (80.0±13.0) | 12.6±0.7 (65.0±9.0) |
| $R_{meas}$ (%) | 14.9±0.7 (113.1±18.5) | 15.0±0.8 (91.9±12.9) |
| Wilson B-factor ($Å^2$) | 19.220±0.457 | 18.994±0.291 |
| Beam divergence (°) | 1.047±0.093 | 1.045±0.091 |
| Mosaicity (°) | 0.130±0.038 | 0.134±0.043 |
| **Refinement statistics** | **$M_{off,n}$** | **$M_{THz,n}$** |
| Resolution range (Å) | 44.57-1.44 (1.48- 1.44) | 44.57-1.44 (1.48- 1.44) |
| Number of refined reflections | 31187±27 (333±3) | 31160±33 (332±4) |
| Number of non-hydrogen atoms | 1877 | 1877 |
| $R_{work}$ (%) | 17.8±0.2 (41.3±1.4) | 17.8±0.2 (42.0±1.4) |
| $R_{free}$ (%) | 20.2±0.3 (39.4±3.0) | 20.3±0.3 (33.8±3.4) |
| RMSD bond angles (°) | 1.675±0.014 | 1.648±0.011 |
| RMSD bond distances (Å) | 0.010±0.000 | 0.010±0.000 |
| Refined B-factor ($Å^2$) | 13.72±0.08 | 13.76±0.07 |

#### Supporting Figures

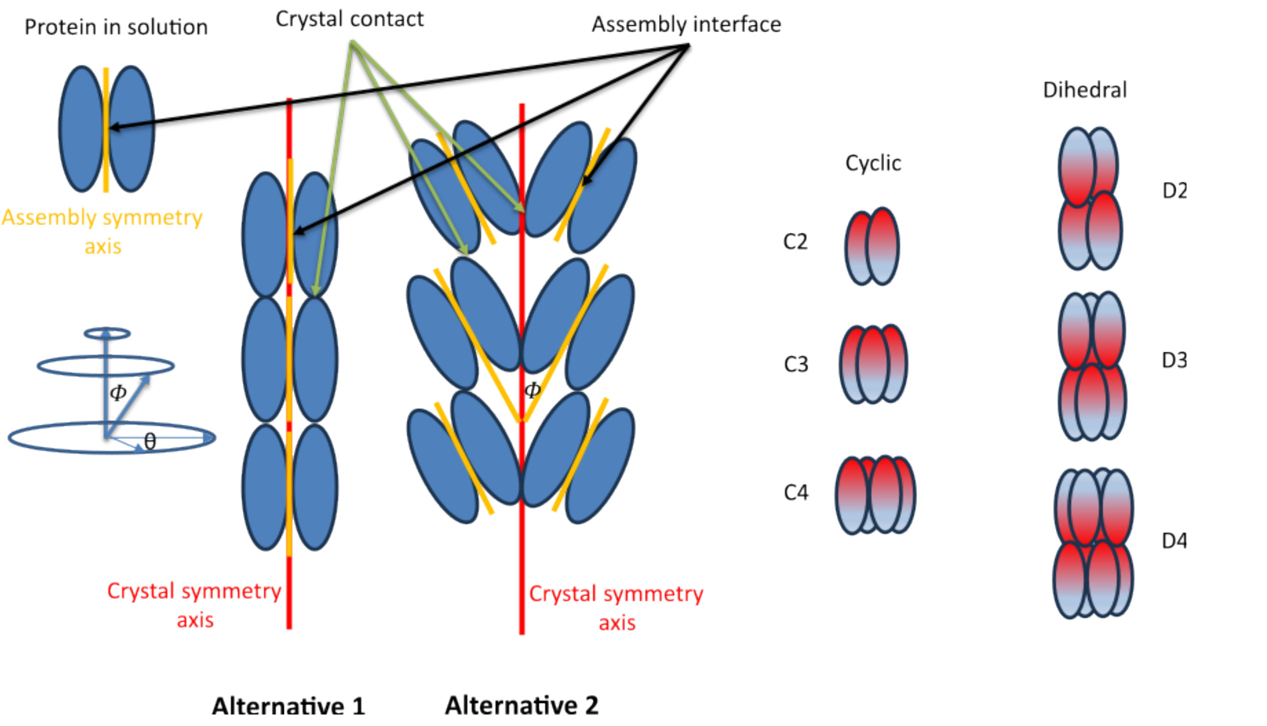


**Figure S1**: Probabilistic description of vector alignment by means of the solid angle, $\Omega = \Omega\,(\Phi, \theta)$. While $\theta$ represents the azimuth, such a probability arises from a bound on the maximum polar angle ($\Phi$) compatible with alignment conditions discernible in experiments (here set to $\overline{\Phi} \approx 1°$). A full solid angle $\Omega = 4\pi$ (unit probability) clearly corresponds to a totally unrestricted $\Phi$. The connection between assembly symmetry axes and crystal symmetry axes is exemplified using a dimer with C2 symmetry. The unique surfaces of the assembly interface and crystal contacts are compared where they deviate from each other. Schematic illustrations depict the cyclic (C2-4) and dihedral (D2-4) point groups utilized in the PDB survey.

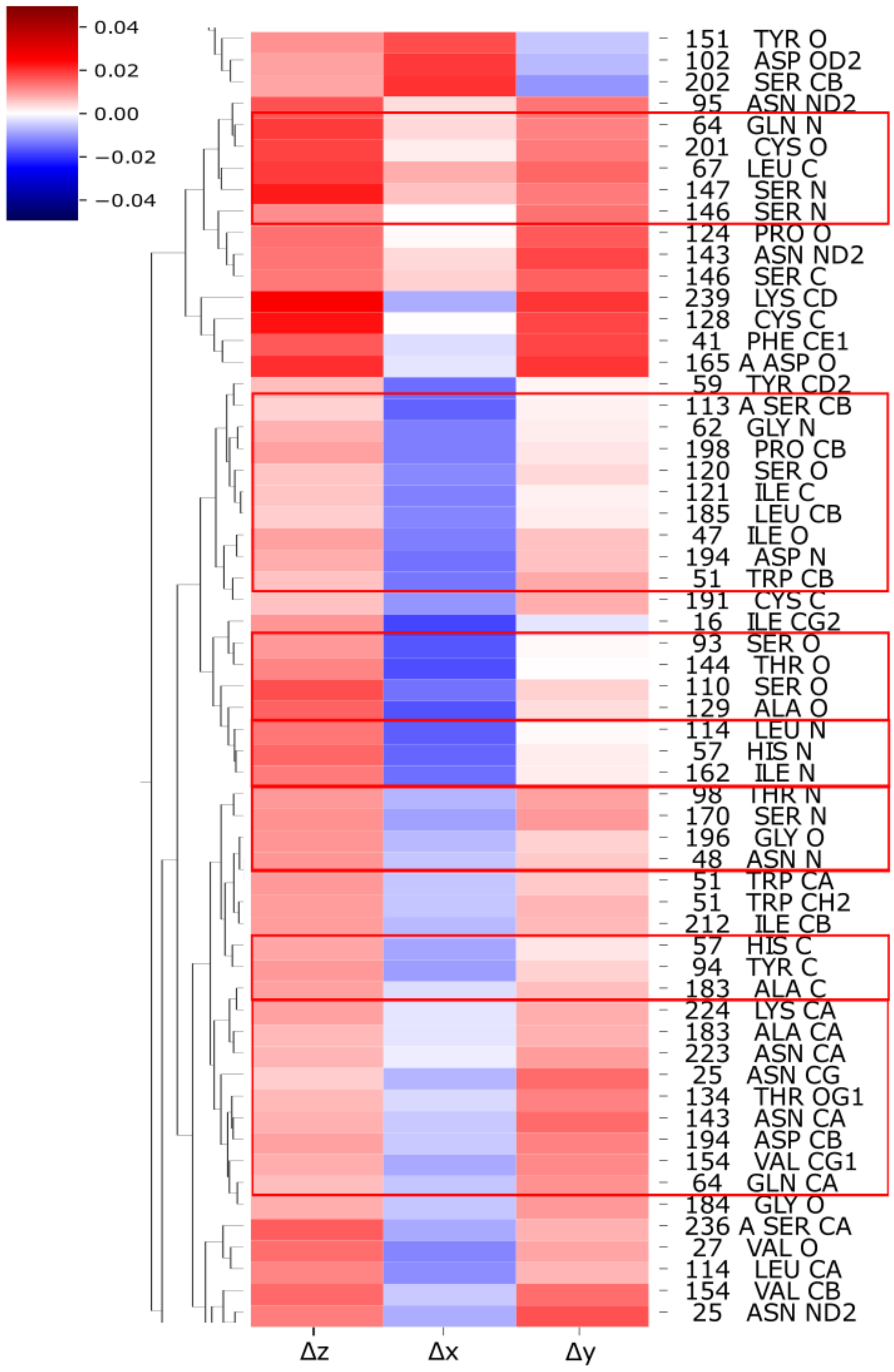


**Figure S2: Enrichment of similar protein atom types among the atoms with most similar** deformation field vectors $\langle M_{THz,n} - M_{off,n} \rangle$ after hierarchical clustering. For better clarity, only 80 out of a total 1629 atoms are displayed.

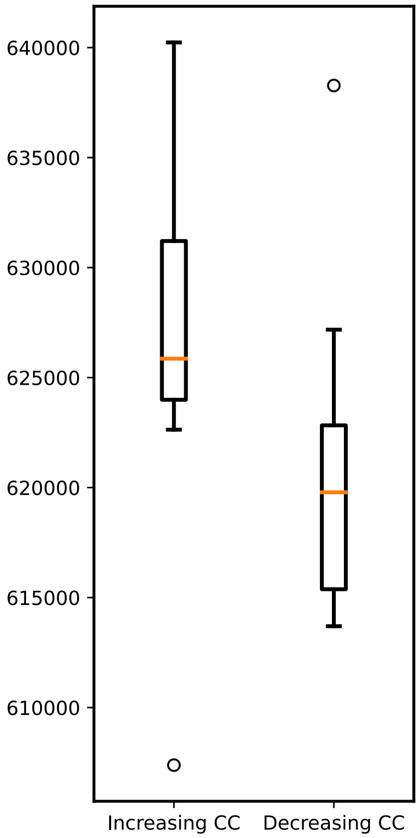


**Figure S3: Numbers of atom pairs in MD simulations with a change in their positional correlation with THz field present versus THz field absent.** Boxplots refer to the numbers of atom pairs that exhibit increased (left) and decreased (right) correlated motions upon introducing a continuous THz electric field in MD simulations. The displayed distributions are based on 10 pairs of MD simulations, with 10 random initial orientations. For each simulated pair, the starting orientation is the same. One is affected by a continuous THz E-field while the other is not.

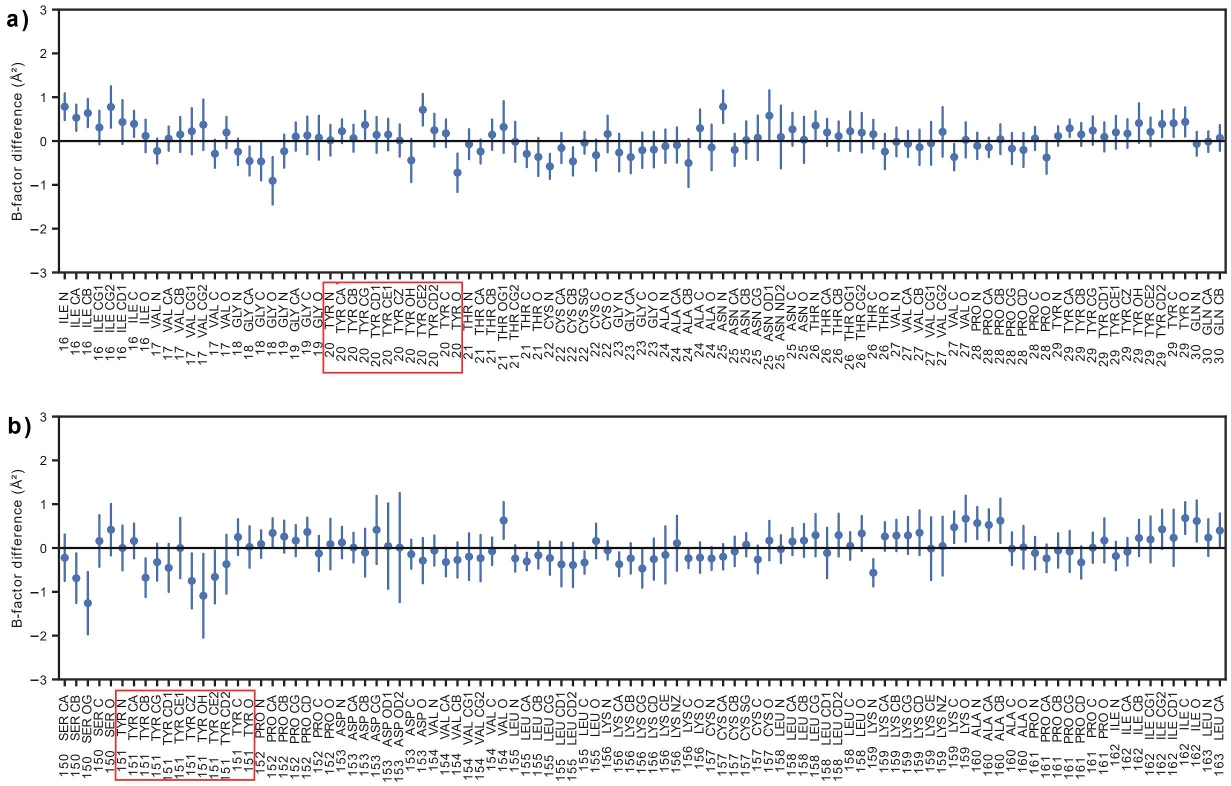


**Figure S4: Deformation intensity distribution**. a) B-factor differences $B_{THz,n} - B_{off,n}$ in the region 16-30 and b) 150-163. The error bars represent the standard deviation.

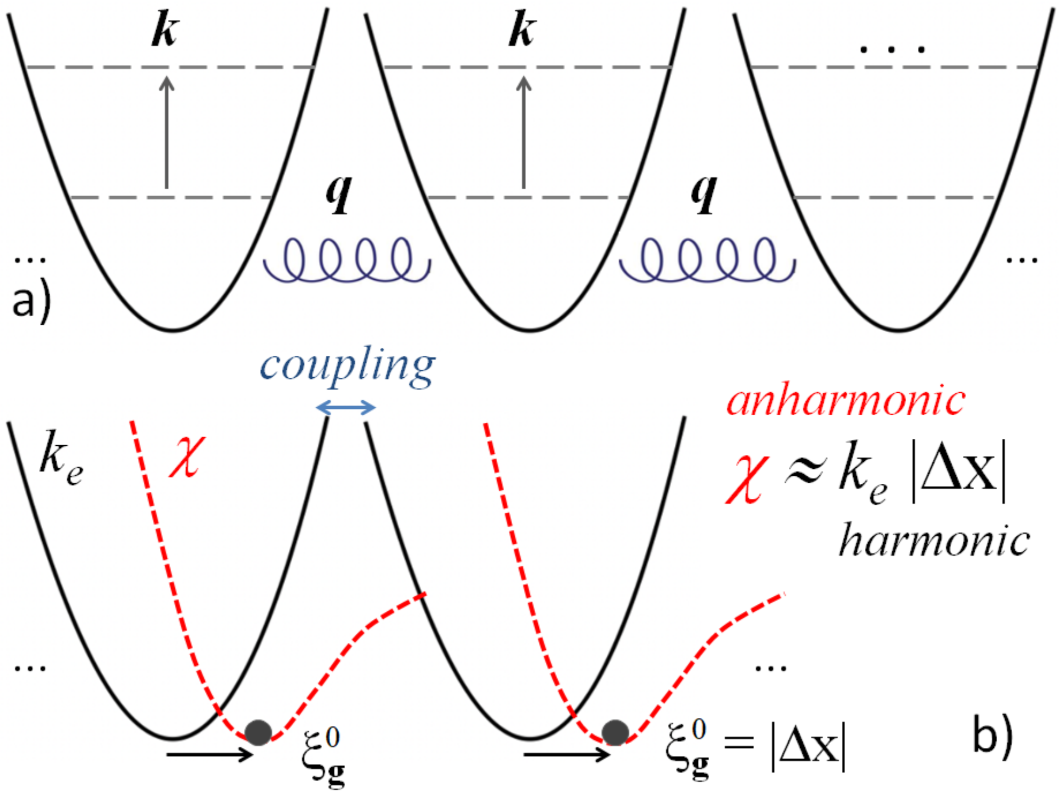


**Figure S5:** Illustrative interpretation of the second quantization scheme (notations are as throughout the work). a) Intramolecular (***k***) and intermolecular (***q***) excitations in their unperturbed state (no coupling). The former, more local in nature, can be assigned to some constituents (dipoles, amino acids, etc.) along the protein chain. The latter are long–range phonon quasiparticles responsible for lattice displacements (or deformations). b) When they couple at some coherent steady state, ***q*** = **g**, an anharmonic interaction develops on lattice atoms (see the coupling term proportional to $\mu_{-\mathbf{g}}S_{(\mathbf{g})}$ on the right side of Eq. S34). As a result, an atom displacement establishes from the balance of harmonic and anharmonic forces acting on the lattice. The displacement $\xi_a$, entraining with the lattice dynamics (Eq. S48), relates analogously with intramolecular modes (***k***) and, for simplicity, is not shown here.

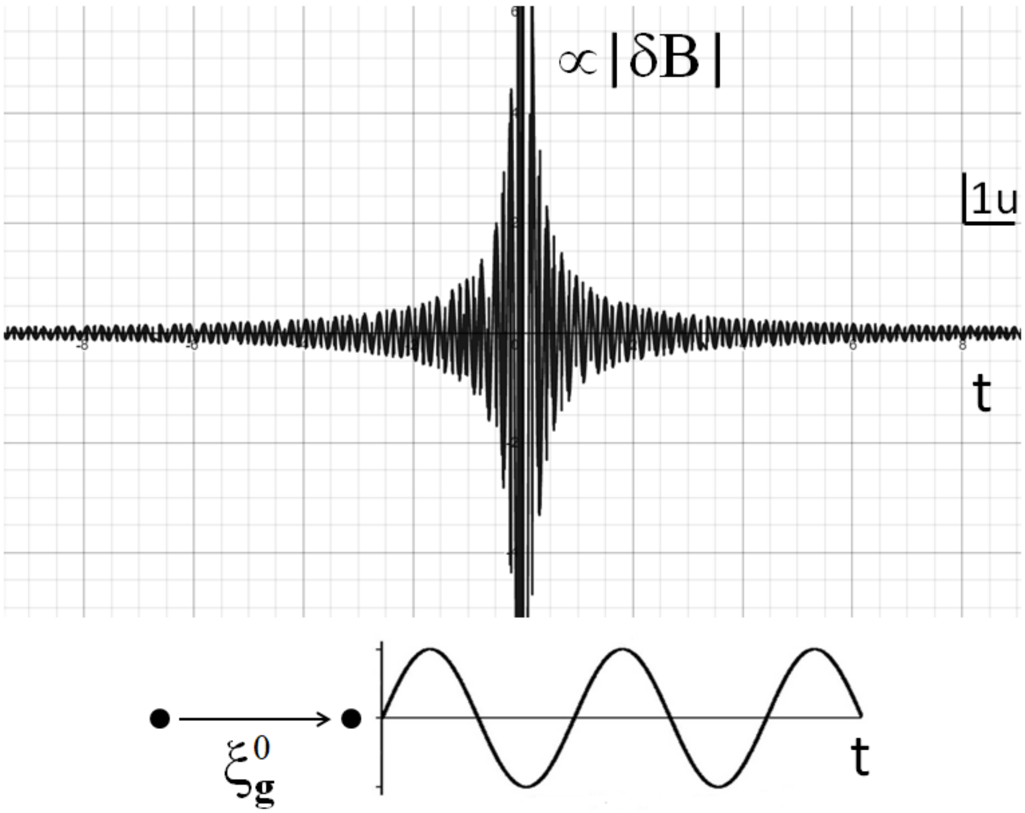


**Figure S6:** Essential scheme for a squeezed coherent state in protein dynamics. Below is a classical wave displacing an atom by a constant lattice shift. Above is the time-dependent part proportional to the B-factor modulation (fluctuations squeeze around the mean atomic positions). It is evaluated from $|\delta B| \propto (1/t)\ [\zeta\Omega_M^{½}\ (4\ \sin(\Omega_M t) - ¼\ \sin(2\Omega_M t)) - \sin(\zeta\Omega_M^{3/2}\ t)]$, for simplicity with $\Omega_M$ = 5 THz and $\xi_g$ = 1 pm ($n_N$ = 1). Symbols denote the same quantities in the main text. Atom displacement and B-factor oscillation are not meant to be on scale.

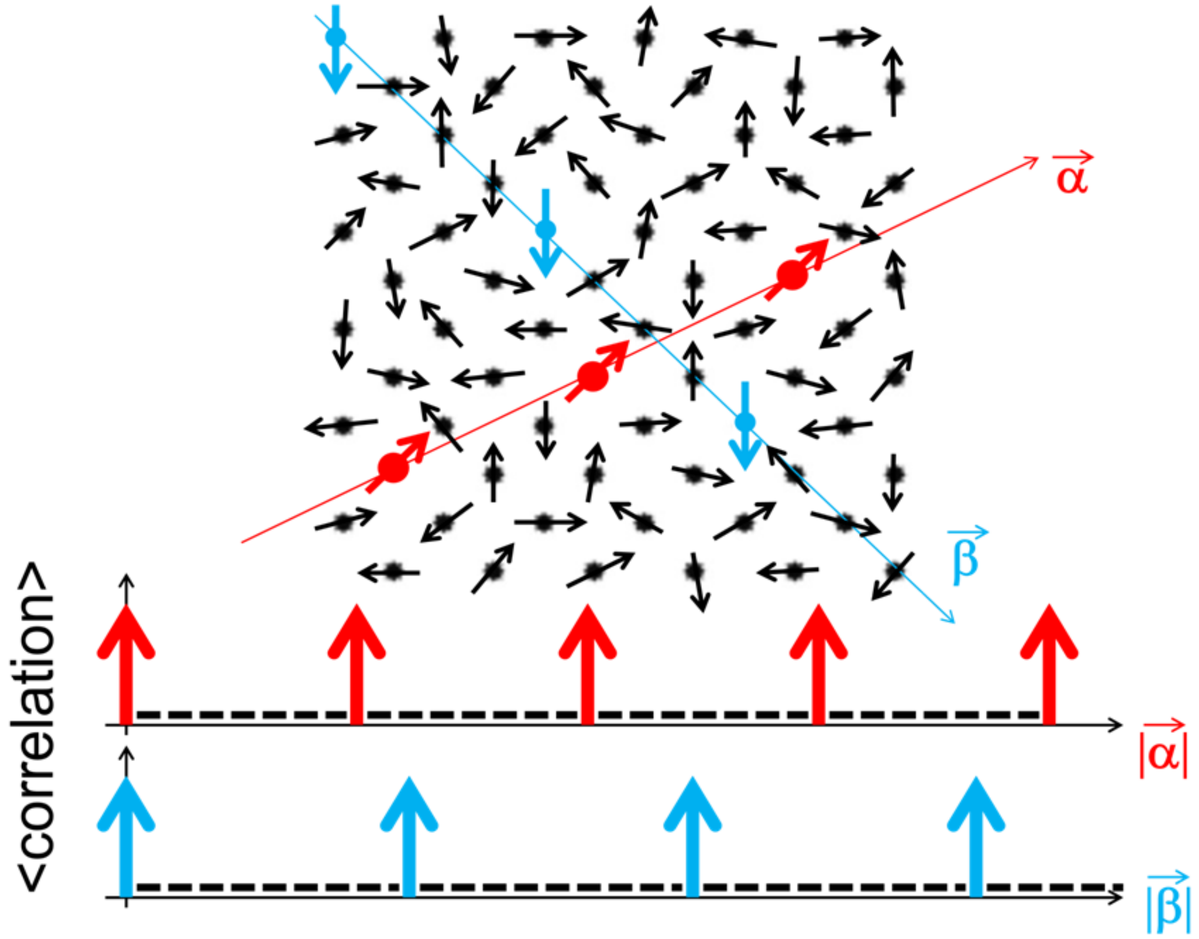


**Figure S7: Idealized scheme for distal correlations in our experiments.** A couple of correlated patterns (red and blue colours) develop in a network of uncorrelated 'spin' units (black spots and arrows) along directions **α** and **β**. Correlation at a given position (k) may be defined as the averaged sum, over the network sites, of pairwise correlations between atom at k and every other atom, to be zero (dashed line) outside **α** and **β**.

**Movie S1: Establishment of a Selective Mode–Locking Phenomenology of Coherent Oscillators**. Illustration of how a mode-locking of displacements between homologous atoms can result into a (partially) coherent structural ordering. Initially, non-bonded tryptophan molecules are randomly oriented, and a dominant displacement of the gamma carbon atom is assumed to be approximately perpendicular to the indole ring. A harmonic movement is animated, starting with a random phase. In the first animation stage, the oscillation directions gradually align to a common wavevector, and phase shifts decrease. In the second stage, steady-state oscillations are repeated, to reveal the rotational symmetry of tryptophan molecules. These types of ordering can be fundamental to the folding of toroid repeat proteins and assembly of protein molecules with cyclic, dihedral and helical symmetry leading to ring-like, propeller-like, barrel-like, and spiral-like structures. Typical examples are microtubules and actin filaments.

If the oscillation is harmonic and unfavorable bonding does not impede rotation, the oscillators can integrate into the entrained system with two opposite orientations and a 180° phase difference while maintaining a positive correlation of displacements along the common wavevector. This is depicted in the third phase of the animation, where randomly oriented tryptophane molecules align their oscillation direction with that of their gamma carbon atom after undergoing a minimal rotation. By careful observation, two distinct groups of tryptophane molecules can be identified, exhibiting twofold symmetry in addition to their rotational symmetry around the common wavevector. These two groups are color-coded as blue and red. In instances where such an oscillator system determines the orientation of protein subunits, it can lead to the formation of stacked rings, such as the octameric stacked ring of Dmc1 with D8 symmetry[56]. In the context of crystallography, a similar relationship exists between space group pairs P622 and P6, or P422 and P4. The oscillators in space groups P6 and P4 possess clear orientations, preventing the joining of an oscillator system with opposite phase, either due to anharmonic oscillation or bonding that hinders head-head and/or tail-tail interactions. On the other hand, P622 and P422 oscillators need to be harmonic and can align with the collective wavevector in both orientations. Line and plane symmetry and the presence or absence of a screw operator, is inconsequential, as they primarily contribute with translation, and the positions of the oscillators do not play a significant role in this model.

The establishment of mode-locking may take millions of cycles. This is in contrast with the accelerated morphing process observed in the transition from a random to an ordered state, which can be easily perturbed, as we have demonstrated in our experiment by letting the protein crystal interact with a THz pulse. However, the dynamic system returns to the same stationary point and begins to restore immediately after the perturbation. It is also realistic to expect that entrained oscillators are not in perfect phase relationship but give rise to a distribution of oscillation directions around the mean orientation of the protein system. Such a picture can take place even at a steady state and different degrees of ordering, depending on the number of oscillators and the bonding/topological constraints of the polypeptide chain.
